# Choosing variant interpretation tools for clinical applications: context matters

**DOI:** 10.1101/2022.02.17.480823

**Authors:** Josu Aguirre, Natàlia Padilla, Selen Özkan, Casandra Riera, Lidia Feliubadaló, Xavier de la Cruz

**Affiliations:** Research Unit in Clinical and Translational Bioinformatics, Vall d’Hebron Institute of Research (VHIR), Universitat Autònoma de Barcelona, P/Vall d’Hebron, 119–129; 08035, Barcelona, Spain; Hereditary Cancer Program, Catalan Institute of Oncology (ICO), ONCOBELL-IDIBELL, CIBERONC, Barcelona, Spain; Institució Catalana de Recerca i Estudis Avançats (ICREA), Barcelona, Spain

**Author notes:** These two authors have contributed equally to this work (Josu Aguirre, Natàlia Padilla). **Corresponding author**; Xavier de la Cruz.

**Keywords:** Clinical variant interpretation, Molecular diagnostics, Cost models, Personalized medicine, *In silico* tools, Pathogenicity prediction

## Abstract

Our inability to solve the Variant Interpretation Problem (VIP) has become a bottleneck in the biomedical/clinical application of Next-Generation Sequencing. This situation has favored the development and use of bioinformatics tools for the VIP. However, choosing the optimal tool for our purposes is difficult because of the high variability of clinical contexts across and within countries.

Here, we introduce the use of cost models as a new approach to compare pathogenicity predictors that considers clinical context. An interesting feature of this approach, absent in standard performance measures, is that it treats pathogenicity predictors as rejection classifiers. These classifiers, commonly found in machine learning applications to healthcare, reject low-confidence predictions. Finally, to explore whether context has any impact on predictor selection, we have developed a computational procedure that solves the problem of comparing an arbitrary number of tools across all possible clinical scenarios.

We illustrate our approach using a set of seventeen pathogenicity predictors for missense variants. Our results show that there is no optimal predictor for all possible clinical scenarios. We also find that considering rejection gives a view of classifiers contrasting with that of standard performance measures. The Python code for comparing pathogenicity predictors across the clinical space using cost models is available to any interested user at: https://github.com/ClinicalTranslationalBioinformatics/clinical_space_partition

**Summaries:** Josu Aguirre earned his doctorate at the Clinical and Translational Bioinformatics group, at the Vall d’Hebron Institute of Research (VHIR).

Natàlia Padilla earned is a post-doctoral researcher at the Clinical and Translational Bioinformatics group, at the Vall d’Hebron Institute of Research (VHIR).

Selen Özkan is a Ph.D. student at the Clinical and Translational Bioinformatics group, at the Vall d’Hebron Institute of Research (VHIR).

Casandra Riera earned her doctorate at the Clinical and Translational Bioinformatics group, at the Vall d’Hebron Institute of Research (VHIR).

Lidia Feliubadalo earned her doctorate at the Universitat de Barcelona, presently she is a high-level technician working at the Catalan Institute of Oncology (ICO) in the diagnosis of hereditary cancers.

Xavier de la Cruz is ICREA Research Professor at the Vall d’Hebron Institute of Research (VHIR). His research interests revolve around the application of machine learning methods to healthcare problems.

## Introduction

The clinical application of Next-Generation Sequencing (NGS) is presently limited by our inability to interpret its results fully [1–4]: we cannot establish, with 100% accuracy, if the sequence variants identified by this technique are benign or pathogenic. This problem, known as the Variant Interpretation Problem (VIP) [1], has important consequences in terms of patient lives and economic cost and is considered one of the challenges determining the future of Genomic Medicine [3].

A promising option to address the VIP is the use of pathogenicity predictors, which are bioinformatics tools that pose the VIP as a classification problem and solve it with machine learning algorithms [5]. Fast and inexpensive, the utilization of these tools is contemplated in the ACMG/AMP [6] guidelines for variant interpretation. However, their large numbers (>50) and moderate accuracies [5] pose a problem when the intended users have to find an adequate tool for their purposes.

To select a pathogenicity predictor for a healthcare application, one should consider how its use affects downstream medical decisions [7]. However, in the description [5] and clinical application [8,9] of predictors, it is common practice to characterize these tools using only performance measures. There are two problems with this practice. First, standard performance measures (e.g., the Area Under the Curve –AUC, or the Matthews Correlation Coefficient –MCC) are blind to the deployment context [7,10]. This limitation would not be serious if the context were constant. However, this is not the case. The diversity of healthcare scenarios is huge [11–14], with variations between and within countries in providers, drug prices, treatment protocols, etc. This diversity can make what is optimal in one scenario become sub-optimal in another. The second problem of using only performance measures for tool selection is that these measures ignore the incomplete coverage of the vast majority of pathogenicity predictors [15]. These tools are reject classifiers [16], that is, classifiers that give no answer to the prediction problem when prediction confidence is low [16]. Although apparently inconsequential, rejection cannot be ignored because there medical actions are associated with it: additional clinical testing, channeling of patients to surveillance programs, etc.

This work addresses these problems in predictor comparison using cost models [16–20]. Cost models are commonly applied in medical decision processes and the assessment of classifiers. In addition to their simplicity and interpretability, a convenient characteristic of cost models is that they include parameters that allow encoding the application context. Here, we use this property to study how context affects tool selection. Our starting point is the cost model derived for reject classifiers [16]. This model combines two opposing terms, one for misclassification or error rate and the other for rejection rate. The misclassification rate treats false positives and false negatives equally, a common approximation [17] that is unrealistic in medical scenarios [18]. Here, we split the misclassification rate into two terms, one for each error type. We then develop the formalism for comparing an arbitrary number of predictors across the range of clinical scenarios. As a reference, we also derive the formalism for the more straightforward case in which rejection is ignored. To illustrate our methodology, we analyze a set of seventeen pathogenicity predictors for missense variants, showing how different methods may be preferred depending on the clinical context.

## METHODS

The cost framework we have created to compare classifiers across the whole space of clinical scenarios is described in the first four sections.

### Cost model for misclassification errors (MISC)

Cost is a measure of the economic and non-economic components that constitute the clinical context of a medical problem [18,21]. These components may be the monetary cost incurred by public institutions, the cost of disease morbidity in undiagnosed patients, etc. For this reason, cost is often utilized to compare clinical tests [18] or to compare machine learning tools for medical applications [16]. Here we adapt the cost framework to the problem of comparing pathogenicity predictors.

In this section, we describe a cost model that only considers misclassification errors [17,19,20]; it does not consider rejection. This model is not the most adequate for the pathogenicity prediction field, where most tools are rejection classifiers. However, because of its simplicity, MISC allows us to introduce the main ideas in the cost-based selection of tools.

Under MISC, c, the average cost of using a pathogenicity predictor in a clinical setting is expressed as:

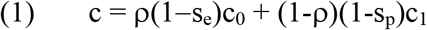

where c_0_ and c_1_ are the misclassification costs: c_0_ is the cost of annotating pathogenic variants as benign, and c_1_ is the cost of annotating benign variants as pathogenic. Each pair (c_0_,c_1_) is characteristic of the clinical setting. ρ and 1-ρ are the probabilities of the two predicted classes, pathogenic and benign variants. ρ is comprised between 0 and 1 and varies with the genome region sequenced and the population of individuals tested. Finally, s_e_ and s_p_ are the sensitivity and specificity of the pathogenicity predictor; they are estimated by testing the predictor in a set of N_p_ pathogenic and N_b_ benign variants as follows:

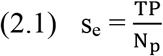

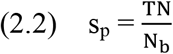

where TP and TN are the numbers of correctly predicted pathogenic and benign variants, respectively.

Following Hernández-Orallo et al. [20], we normalize c using c_T_ = c_0_+c_1_, the cost magnitude. We obtain a normalized average cost, rc:

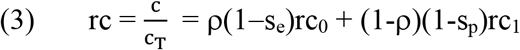

where rc_0_=c_0_/c_T_ and rc_1_=c_1_/c_T_, and rc_0_+rc_1_=1.

Working with rc simplifies subsequent analyses because c_0_ and c_1_ have an indefinite variation range, while their normalized equivalents, rc_0_ and rc_1_, vary between 0 and 1. It must be noted that comparing predictors with either c or rc gives the same result because both values are monotonically related.

We simplify equation (3) by replacing rc_0_ with 1-rc_1_:

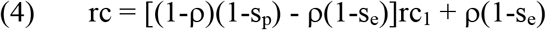

In equation (4), clinical scenarios are represented by their rc_1_ values. The range of rc_1_ values, the interval **I**=(0,1), is that of all possible clinical scenarios, and we will call it ‘clinical space’.

### Predictor comparison across clinical scenarios with the MISC model

For a given clinical scenario (defined by its rc_1_ value), it is easy to use rc to compare N predictors; we only need to compute and sort their respective rc values. The method of choice will then be the one with the lowest rc.

Here we want to generalize the comparison of N predictors to all possible scenarios (all rc_1_ values) because nothing guarantees that the same method will have the lowest rc. That is, we are looking for a division of **I** into a set of sub-intervals such that only one method per interval has the lowest rc and this method is different between intervals.

The protocol we present is based on the fact that we can interpret the cost equation (4) for each predictor as a line in rc_1_. Comparing predictors in terms of cost is equivalent to finding the rc_1_ value at the intersection between their lines. This value will divide **I** into two parts with one prevalent predictor in each. If the rc_1_ at the intersection falls outside **I**, then only one method will prevail over the whole interval. The extension of this idea to N predictors is as follows.

For a set {M_i_, i=1,N} of N predictors, the desired division of **I** is obtained by executing the following four-step protocol:

**Step 1.** Solve in rc_1_ all the equations rc(M_i_)=rc(M_j_), (1≤i≤N-1; i<j≤N). The set of solutions obtained is: 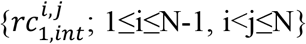, where the indexes i and j refer to the M_i_-M_j_ comparison.

**Step 2.** Eliminate from the set of solutions all the points falling outside **I**. Then, sort the remaining values: 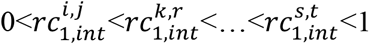. Note that between two rc_1,int_ adjacent values, there is no pair of rc lines crossing each other.

**Step 3.** For each of the associated intervals 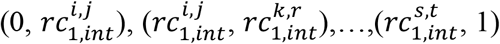, find the predictor with the lowest rc at the interval’s midpoint. This predictor will have the lowest rc value all over the chosen interval because, within intervals, rc lines do not cross each other (see Step 2).

**Step 4.** Unify those adjacent intervals for which the same predictor has the lowest rc, repeating this step until all adjacent intervals correspond to different methods. Because of this univocal correspondence between intervals and predictors, the resulting list of intervals, {**I**_i_=(a_i_,b_i_); 1≤i≤m} gives the desired distribution of predictors across **I**. Note: m<N, since not all the methods are necessarily present in the final list.

#### Interpreting the solution

The list of intervals **I**_i_ and their associated predictors are the answer to the problem of comparing N predictors across the clinical space with the MISC model. A simplified version of this result is obtained by computing the size of each interval, which is equal to |**I**_i_|= b_i_-a_i_, for **I**_i_=(a_i_,b_i_). This number is the fraction of clinical scenarios where Mi is preferable to the other predictors.

It must be noted that the list of intervals/predictors obtained depends on the value of ρ. This dependence will be explored when applying this formalism to a set of seventeen chosen pathogenicity predictors (Supplementary Table 1).

A python implementation of this procedure, CSP-norej (Clinical Space Partition, no rejection), is available at: https://github.com/ClinicalTranslationalBioinformatics/clinical_space_partition

### Cost model for misclassification errors plus rejection (MISC+REJ)

Here, our starting point is the cost model for reject classifiers [16], which we extend, replacing the misclassification error with the more general expression in equation (1). Then, for a given pathogenicity predictor, c, the average cost of using this tool in a clinical setting is expressed as:

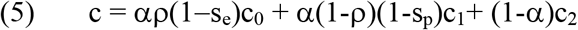

where some parameters, s_e_, s_p_, ρ, c_0_, and c_1_, are the same as in equation (1). The two new parameters are: c_2_, the cost associated with rejection; and a, the coverage of the predictor. The latter is directly related to the rejection rate, equal to (1-a). We use a in equation (5) because it is a descriptor commonly employed in the pathogenicity prediction field [15]. It is computed as:

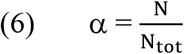

where N is the number of cases from a total of N_tot_ variants (a mixture of pathogenic and benign cases) for which the predictor generates a result.

As before, instead of c we will use rc, the normalized average cost, obtained after dividing both sides of equation (5) by c_T_ (=c_0_+c_1_+c_2_):

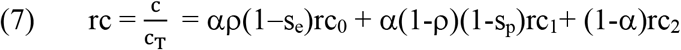

where rc_i_=c_i_/c_T_ (i=0,2) are comprised between 0 and 1, and rc_0_+rc_1_+rc_2_=1. As for MISC, comparing predictors with either c or rc gives the same result because both quantities are monotonically related (equation (7)).

We reduce the number of parameters in rc by replacing rc_2_ with 1-rc_0_-rc_1_ in (7):

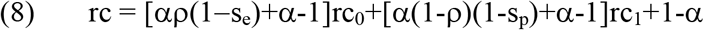

rc is now defined over a triangular region **T** in the rc_0_-rc_1_ plane, bounded by the axes rc_0_, rc_1_, and the line rc_0_+rc_1_=1. **T** is conceptually equivalent to **I** in the MISC case: each point in **T** corresponds to a clinical scenario. We will refer to **T** as ‘clinical space’ also. However, **I** and **T** differ in that the second is two-dimensional; i.e., clinical scenarios are represented by (rc_0_, rc_1_) pairs, not by a single value.

In Supplementary Materials, Appendix 1, we provide an extended description of the methodology described below, including proofs for the most relevant results.

### Predictor comparison across clinical scenarios with the MISC+REJ model

The comparison of N predictors in a clinical scenario defined by a pair (rc_0_, rc_1_) follows the same steps as before: we only have to compute and sort their rc values. The method of choice will be the one with the lowest rc.

However, extending the comparison to all possible clinical scenarios is now more difficult because we have two parameters, rc_0_ and rc_1_, instead of one. We have to divide a two-dimensional figure, instead of a one-dimensional interval, into a set of m regions r_k_ (k=1,m), within each of which a single method prevails over the others. To explain how we obtain these regions, we describe the case of two predictors (N=2), and then extend it to an arbitrary number of predictors.

Let M_i_ and M_j_ be two pathogenicity predictors, and rc(M_i_) and rc(M_j_) their respective rc’s. We seek a division of **T** into two regions: r_i_, where M_i_ is preferable to M_j_ (rc(M_i_) < rc(M_j_)), and r_j_, where the opposite is the case (rc(M_i_) > rc(M_j_)). The boundary between r_i_ and r_j_ is defined by the equation rc(M_i_)=rc(M_j_), which, using equation (8) for rc(M_i_) and rc(M_j_), gives:

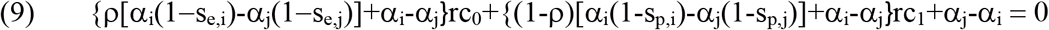

where s_e,k_, s_p,k_, and α_k_ are the sensitivity, specificity, and coverage of predictor M_k_ (k=i,j). Equation (9) shows that the boundary sought is a line (Figure 3a), which we will call l_ij_, in the rc_0_-rc_1_ plane.

When l_ij_ crosses **T**, it divides it into two convex polygons (Figure 3a), corresponding to the r_i_ and r_j_ regions. If l_ij_ does not cross **T**, then only one of the two methods will have the lowest rc in all **T** points.

From Equation (9), we see that l_ij_ depends on ρ; consequently, different values of this parameter may change r_i_ and r_j_. This dependence will be explored below, when applying this formalism to a set of seventeen chosen pathogenicity predictors (Supplementary Table 1).

The generalization of this procedure to more than two predictors is based on the following. First, the lines associated to all possible pair comparisons between predictors divide **T** into a set of convex polygons (P_N_), within each of which a single predictor prevails. Proof of this result is available in the Supplementary Materials (Appendix 1, Section 2). Second, grouping these polygons according to their associated methods gives the desired regions. And third, these polygons can be found using an adapted version of the Breadth First Search algorithm. Below we describe in more detail the two last points.

#### Finding the r_k_ (k=1, m) regions from the polygons in P_N_

The polygons in P_N_ can be used to obtain the r_k_ regions as follows. First, we find the predictor with the lowest rc within each polygon. To this end, for each polygon (i) we compute the average of its vertices; (ii) we compute the rc value for each predictor at this average point; and (iii) we sort the resulting rc’s and choose the method with the lowest rc. This method will be the one associated to the polygon. At the end of this procedure, we have a list of polygons and their associated methods. Now, we only need to merge the polygons associated with the same predictor to obtain the desired *r_k_* regions. For example, if in P_N_ there are three polygons associated with the predictor PolyPhen-2, merging them will give a region associated with PolyPhen-2. Note that m, the number of regions, is not necessarily equal to that of predictors.

#### Using an adapted Breadth First Search (BFS) to generate polygons in P_N_

Knowing the polygons in PN is required to find the r_k_ (k=1,m) regions. To generate these polygons we will first identify their vertices. They are the intersection points between the lines l_ij_ and between these lines and the sides of the triangle. Now, we can loop over these points, enumerating the polygons meeting at each of them. We can model this part as a cycle enumeration problem in graph theory.

Our starting point is the unweighted, undirected graph G(V, E), whose set of vertices, V, and edges, E, correspond to VP and EP, the sets of vertices and edges of the polygons, respectively. Because the list of vertices of a polygon is formally equivalent to that of a cycle, we can reformulate the original looping through VP elements as looping through V elements. For each v_i_ ∈ V, we will use BFS as shortest cycle generator. Because, in some instances, the resulting cycles correspond to figures with unwanted geometrical properties, we will keep only those cases that meet seven conditions (C1-C7, see Supplementary Material, Appendix 1, Section 3) designed to ensure that the associated figures correspond to polygons in P_N_. It can be shown that the exhaustive application of this combination of BFS and C1-C7 when looping through the vertices in G(V, E) produces the list of polygons in P_N_. The proofs of all the lemmas and propositions behind this procedure are given in the Supplementary Material (Appendix 1, Section 3).

##### Interpreting the solution

The r_k_ regions and their associated predictors are the answer to the problem of comparing N predictors across the clinical space with the MISC+REJ model. A simpler version of this result is obtained by computing each r_k_ region’s surface. This number is the fraction of clinical scenarios where the predictor associated to r_k_ is preferable to the other predictors, in terms of rc.

A python implementation of this procedure, CSP-rej (Clinical Space Partition, rejection), is available at: https://github.com/ClinicalTranslationalBioinformatics/clinical_space_partition

We want to mention that the code reproduces the results presented in this work and allows users to explore other combinations of predictors. However, it is known that geometric computations may present robustness problems [22], which in our case can happen for low ρ values or large numbers of predictors. Therefore, we cannot discard the possibility that the program sometimes fails to find the division of **T**. In these cases, the program will issue an error message.

### Variant dataset

For each predictor, we estimated the three performance parameters used for cost models, sensitivity (s_e_), specificity (s_p_), and coverage (α), in a set of benign and pathogenic variants retrieved from the dbNSFP database [23], version 4.0a, release: May 3, 2019. This version was chosen because it was released after the predictors’ publication dates, thus preventing the effect of first-order circularities [24] in the estimate of sensitivities, specificities, and coverages. There is only one exception to this rule, the predictor EVE [25], published in 2021. However, because this method is unsupervised it is immune to circularity problems.

We imposed three filters on the variants retrieved. First, they should not affect a splicing site. Second, the ClinVar Review Status of the variants should be: ‘Practice guideline’, or ‘Expert Panel’ or ‘Criteria provided, multiple submitters, no conflicts’. And third, we unified the clinical significance classes as follows: ‘Benign’ and ‘Likely_benign’ variants were labeled as ‘benign’; ‘Pathogenic’ and ‘Likely_pathogenic’ variants were labeled as ‘pathogenic’. The resulting dataset comprised 1902 variants, 809 pathogenic and 1093 benign, from 903 proteins.

### Pathogenicity predictors

We have used seventeen pathogenicity predictors: PolyPhen2-HDIV [26], PolyPhen2-HVAR [26], SIFT [27], CADD [28], MutationTaster2 [29], MutationAssessor [30], REVEL [31], LRT [32], PROVEAN [33], MetaLR [34], MetaSVM [34], VEST4 [35], MutPred [36], PON-P2 [37], SNAP2 [38], EVE [25], and PMut [39]. For each variant in our dataset, we retrieved the pathogenicity prediction of these tools from the dbNSFP [23] database, except for PON-P2, SNAP2, and PMut, for which we used the corresponding website.

The methods were chosen to represent the range of sensitivities, specificities, and coverages in this problem.

## RESULTS

### Application of the MISC cost model to the comparison of pathogenicity predictors across clinical scenarios

In MISC, each pathogenicity predictor is characterized by a line relating normalized cost (rc) and clinical scenario (represented by its rc_1_ value) (Equation (4)). Figure 1a shows the lines for the seventeen predictors analyzed for ρ=0.5.

**Figure 1.**
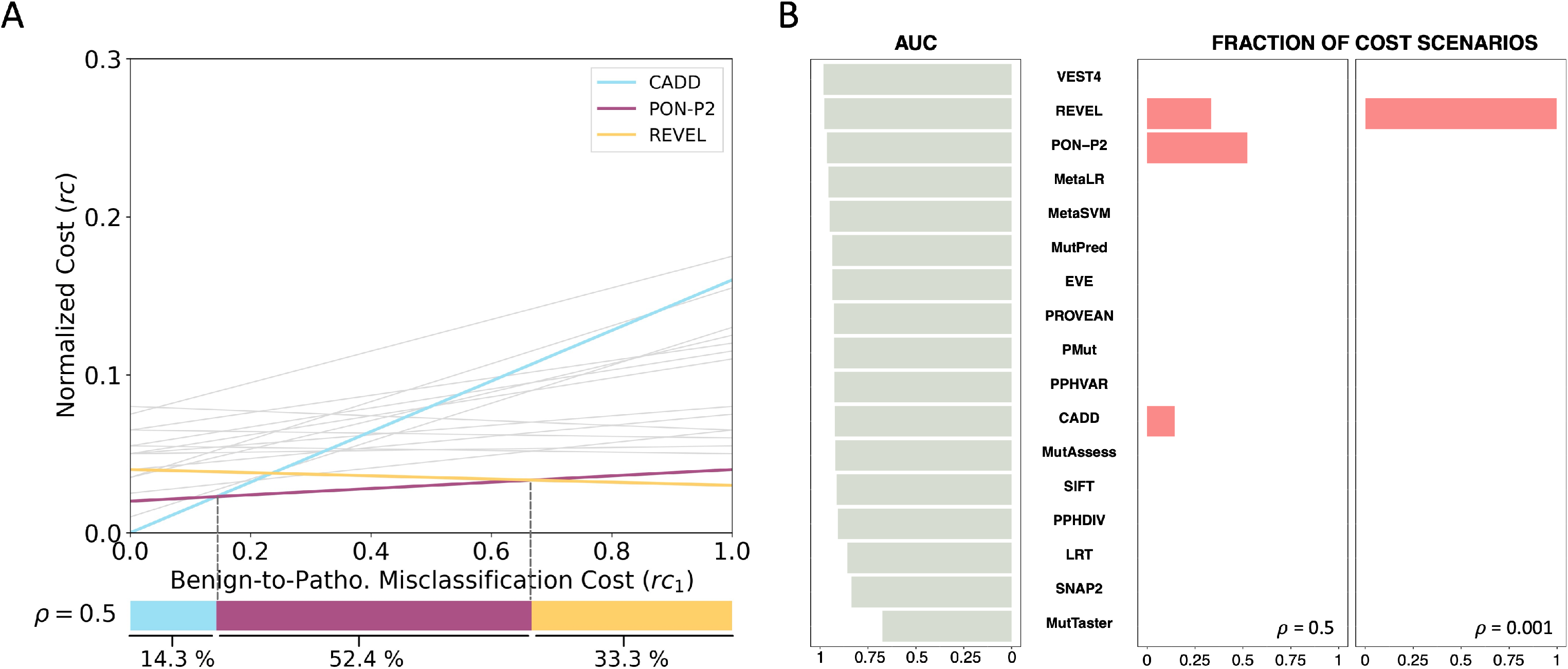
Application of the cost model MISC to seventeen pathogenicity predictors. **a,** The figure describes the partition of the clinical space between predictors under the MISC model. Each predictor is represented by a line whose points correspond to the normalized cost of the predictor (rc, y-axis) at a given value of the benign-to-pathogenic misclassification cost (rc_1_, x-axis). All the lines are grey, except those corresponding to PON-P2 (purple), REVEL (yellow), and CADD (blue), the only methods that prevail over the others, in terms of cost, in a region of the clinical space. These methods and their associated regions are found using the procedure described in the Methods section ‘Predictor comparison … with the MISC model’. The size and location of the regions are shown at the bottom of the figure, in a bar representing the clinical space. Each color corresponds to one of the chosen methods. **b**, The figure shows the contrasting views on predictors provided by AUC (grey bars, left side) and cost models (pink bars, right side; results for ρ=0.5 and ρ=0.001). The meaning of the bars is different. The grey bars indicate the AUC value of a method. The pink bars indicate the fraction of scenarios where the method prevails over the others in terms of cost. If we chose our preferred method using AUC, we would select VEST4. However, for both values of ρ, this method is not cost-effective in any region of the clinical space (VEST4 has no pink bar). We see that PON-P2, which is the third candidate according to AUC, is the most cost-effective method in 52% (size of the pink bar) clinical scenarios. The contrast is stronger for CADD, which ranks eleventh for AUC but is the most cost-effective method in over 14% of scenarios when ρ=0.5.

For a given rc_1_, the line with the lowest position (lowest rc value) corresponds to the preferred predictor. To extend this comparison to all possible clinical scenarios (rc_1_ values) we use the procedure described in the Methods section ‘Predictor comparison … with the MISC model’. We find that the clinical space can be divided into three sub-intervals. We show these sub-intervals at the bottom of Figure 1a, in the form of three adjacent colored bars whose sizes reflect the fraction of clinical scenarios where the associated methods (CADD, PON-P2, and REVEL) are preferred. Above each interval, we see that the line of the associated predictor occupies the lowest position compared to the other lines. In summary, the predictors CADD, PON-P2, and REVEL are the solution to the problem of selecting the preferred tools across the clinical space.

One noteworthy aspect of this result is that there is no single cost-optimal method across clinical contexts; that is, the tool of choice depends on the clinical scenario. This is in contrast to the standard strategy for choosing pathogenicity predictors, where the top-ranking tool according to some performance measure (e.g., AUC, MCC, etc.) is always preferred. That is, a unique method is chosen, regardless of the application scenario. To illustrate this point, we sorted the seventeen predictors according to their AUC (Figure 1b), finding that VEST4 was the top-ranking predictor and, according to this criterion, it would be the tool of choice. Interestingly, from the point of view of cost, there is no scenario where VEST4 is preferable to other predictors. A better coincidence is obtained using MCC instead of AUC, but the general problem remains (Supplementary Figure 1a).

To close this section, we want to comment on the effect of ρ, the frequency of the pathogenic variants, on the cost-based tool selection problem. As part of the cost formalism, this parameter will affect the election of the preferred predictor. For example, if we change ρ from 0.5 to 0.001, we see (Figure 1b) that REVEL prevails over the other methods, predominating in 99.8% of the cost scenarios. When presenting the results of the MISC model, we have used ρ=0.5. This value reflects the fact that the patient population is a biased sample of the general population. Therefore, we expect the fraction of pathogenic variants to be closer to that of benign variants. However, other values may be chosen for other applications. For example, low ρ’s may be preferable for population screenings. In Figure 2, we explore the effect of different ρ’s on the partition of the clinical space between methods. We observe a transition between REVEL and PON-P2 as we go from low to high ρ’s, reflecting the higher sensitivity of PON-P2 (0.96) relative to REVEL (0.92).

**Figure 2.**
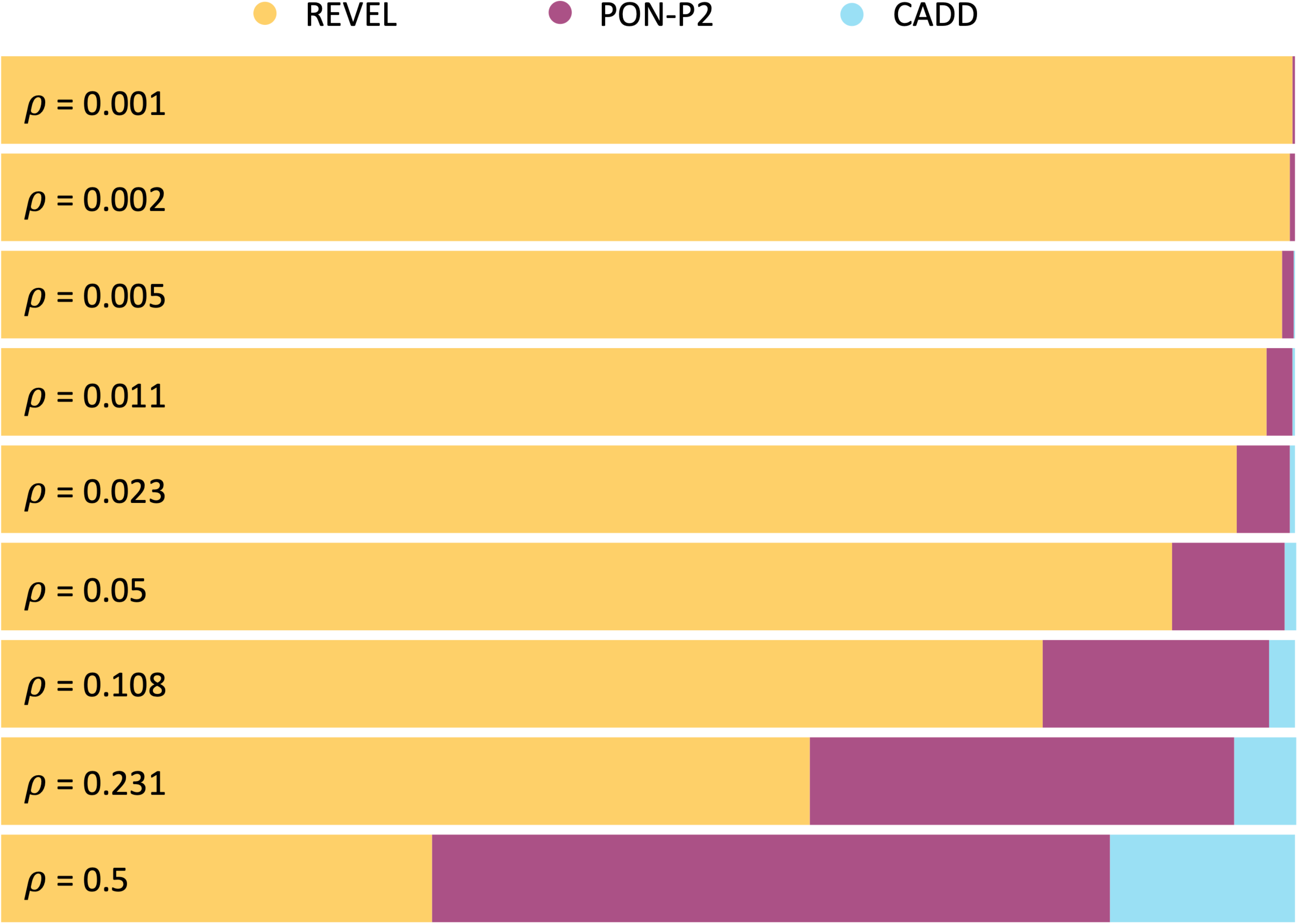
Dependency on ρ of the distribution of pathogenicity predictors in the clinical space, under the MISC model. The figure shows that different values of ρ induce different divisions of the clinical space between predictors. Each horizontal bar represents one of these divisions for a given ρ. The different colors in the bar indicate the size and location of the regions where a single predictor prevails over the remaining sixteen. Only three predictors (REVEL, PON-P2, CADD) appear almost always: the remaining tools are not cost-optimal in any region of the clinical space under the MISC cost model.

### Application of the MISC+REJ cost model to the comparison of pathogenicity predictors across clinical scenarios

Here, we illustrate the use of MISC+REJ for the set of seventeen predictors (see Methods). In this model, predictors are represented by planes (Equation (8)) whose relative positions define the regions where a single predictor prevails over the others. Threedimensional representations involving multiple planes are difficult to analyze. For this reason, we eliminate the rc axis from our plots to represent our results and use only the clinical space **T** (a triangle in the rc_0_-rc_1_ plane, see Figure 3a). This decision does not affect the interpretation of the results, which only requires visualization of the r_k_ regions.

**Figure 3.**
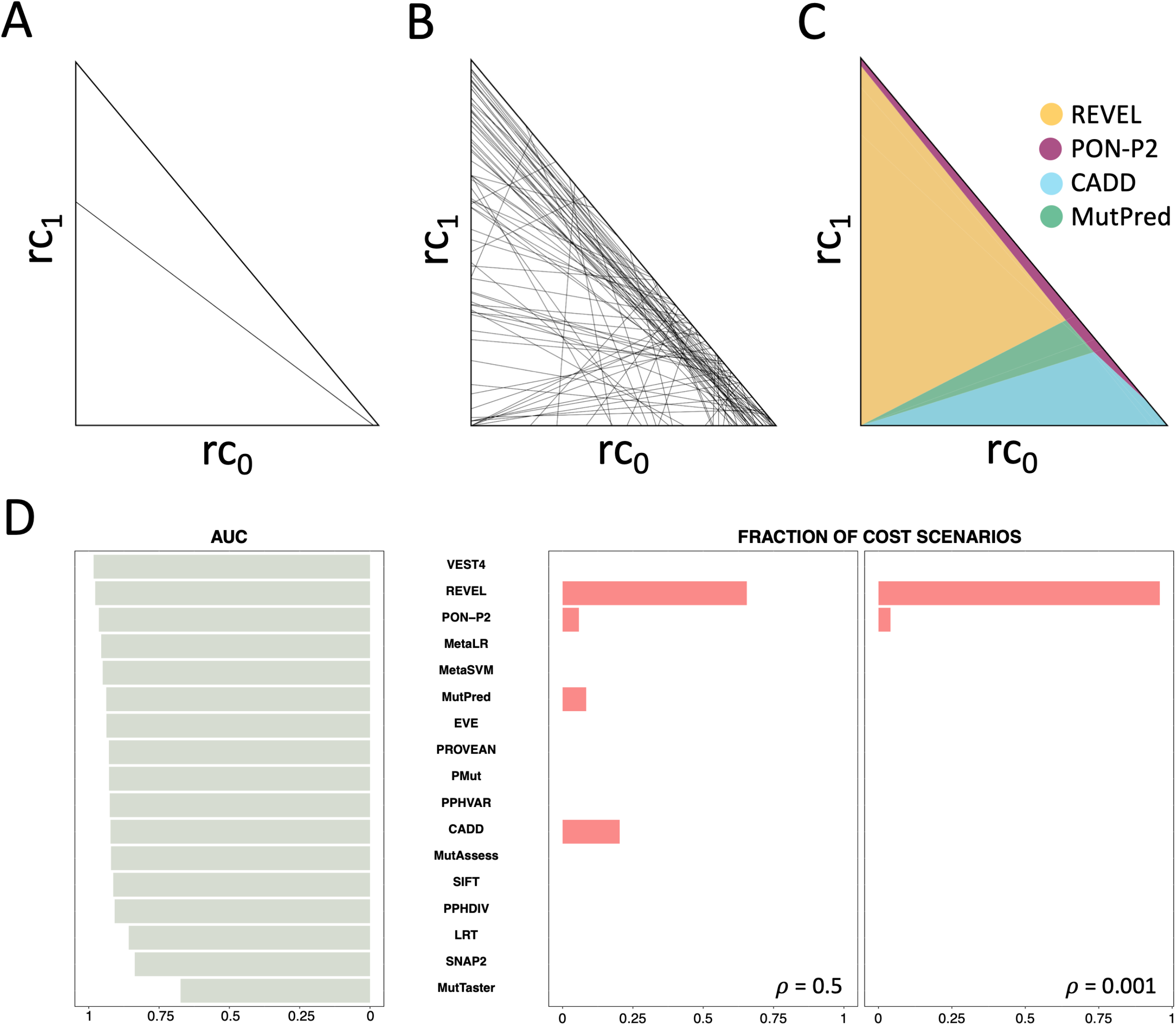
Application of the cost model MISC+REJ (misclassification errors+rejection) to seventeen pathogenicity predictors. The figure describes the partition of the clinical space, **T**, between predictors, under the MISC+REJ model. These methods and their associated regions are found using the procedure described in the Methods section ‘Predictor comparison … with the MISC+REJ model’. In (**a**) we show the comparison of two predictors. Under the MISC+REJ, the clinical space is a triangle. The line traversing the triangle results from comparing two predictors (Equation (9)). It divides the triangle into two polygonal regions within which a single method is cost-optimal. **b**, When all the possible pair comparisons are made, the corresponding lines form a complex pattern of polygons; inside each polygon, a single predictor is cost-optimal. **c,** Finally, we computationally find and unify these polygons using an adapted version of the Breadth-first Search algorithm. The resulting regions, shown with different colors, indicate those parts of the clinical space where a given predictor prevails from the cost point of view. **d,** The figure shows the contrasting views on predictors provided by AUC (grey bars, left side) and cost models (pink bars, right side; results for ρ=0.5 and ρ=0.001). The meaning of the bars is different. The grey bars indicate the AUC value of a method. The pink bars indicate the fraction of scenarios where the method prevails over the others in terms of cost. If we chose our preferred method using AUC, we would select VEST4. However, for both values of ρ, this method is not costeffective in any region of the clinical space (VEST4 has no pink bar). We see that REVEL, the second candidate according to AUC, is the most cost-effective method in many clinical scenarios (long pink bar). The contrast is stronger for CADD, which ranks eleventh for AUC but is the most cost-effective method in over 20% of scenarios, when ρ=0.5.

For a given pair (rc_0_, rc_1_), the method with the lowest rc value (Equation (8)) corresponds to the preferred predictor. To extend this comparison to all possible clinical scenarios ((rc_0_, rc_1_) pairs) we use the procedure described in the Methods section ‘Predictor comparison … with the MISC+REJ model’. This procedure divides the clinical space into a mosaic of polygons (Figure 3b) that is processed to give four regions (Figure 3c), corresponding to the predictors REVEL, CADD, MutPred, and PON-P2. The surface of these regions is proportional to the fraction of clinical scenarios where the associated methods prevail. The results are obtained for ρ=0.5.

As before, there is no single cost-optimal method across clinical contexts. There are, however, some remarkable differences in the distribution of the predictors. Apart from the presence of a new method, MutPred, we observe an important change for the remaining three. More precisely, REVEL now prevails in more scenarios than PON-P2. The rejection rate plays an essential role in this reversal: both predictors have similar sensitivities and specificities (see Supplementary Table 1), but REVEL has a 0% rejection rate, while that of PON-P2 is 54%.

These results confirm that under MISC+REJ the tool of choice depends on the clinical scenario. Again, this view contrasts with the static picture conveyed by performance measures. For example, a comparison with the AUC ranking of the predictors (Figure 3d) shows that VEST4, the predictor with top AUC, does not prevail in any clinical scenario. After excluding VEST4, the coincidence improves: the second predictor classified, REVEL, is the one associated with more scenarios (Figure 3c). However, CADD, which prevails in 20% scenarios, ranks eleventh with AUC (Figure 3d). Repeating the analysis with MCC (Supplementary Figure 1b) shows that this parameter does not capture the REVEL-PON-P2 reversal observed with cost models. The result for CADD is similar to that for AUC.

To close this section, we explore the impact of ρ on the tool selection problem under the MISC+REJ model. In Figure 3d, we see that using ρ=0.001 instead of ρ=0.5 does not substantially alter the results for REVEL, which prevails over most clinical scenarios. However, CADD and MutPred disappear. Looking at the results for different ρ’s (Figure 4), we find a clear predominance of REVEL. This predominance has several origins. One is the null rejection rate of REVEL relative to PON-P2. The second is its better specificity (0.94) relative to CADD (0.68) and MutationTaster2 (0.87). This effect becomes clear when we see how the surface occupied by CADD and MutationTaster2 grows as the fraction of benign variants grows, i.e., as ρ goes from 0.01 to 0.5.

**Figure 4.**
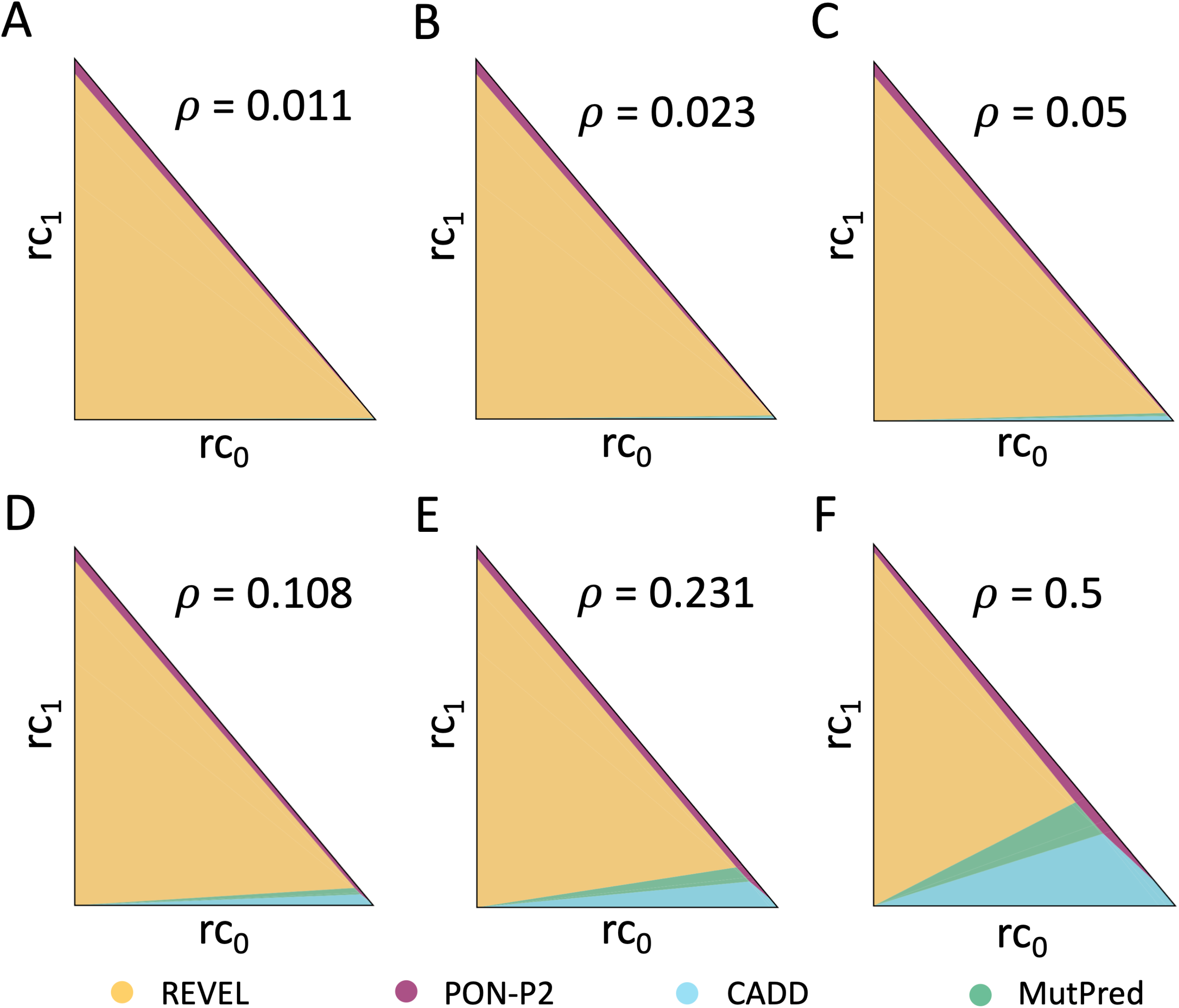
Dependency on ρ of the distribution of pathogenicity predictors in the clinical space, under the MISC+REJ model. Each triangle represents the clinical space for an ρ value. **a,** ρ=0.011; **b,** ρ=0.023; **c,** ρ=0.05; **d,** ρ=0.108; **e,** ρ=0.231; **f,** ρ=0.5. We chose ρ values such that each order of magnitude is evenly sampled. The colored regions correspond to the cost scenarios where the same predictor is cost-optimal. Only four predictors (REVEL, PON-P2, CADD, MutPred) always appear: the remaining tools are not cost-optimal in any region of the clinical space for any ρ, under the MISC+REJ cost model.

## DISCUSSION

In this work, we adapt cost models [16–20] to the problem of selecting pathogenicity predictors in a multi-scenario context. Our proposal constitutes an advance relative to the use of standard performance measures, like AUC or MCC, that give a view of classifiers inaccurate for deployment processes [7]. This is because these measures do not incorporate context descriptions in their definitions; consequently, they give the same ranking of methods regardless of the clinical context. Additionally, each measure has its problems. Several limitations of AUC are known [7,40], e.g., its averaging over sensitivity/specificity trade-offs which may be problematic in clinical applications. For MCC, its symmetry in the sensitivity and specificity [41] means that predictors with opposite sensitivity/specificity trade-offs have the same MCC, an undesirable feature for clinical applications. Another limitation of these measures is that they ignore an key aspect of pathogenicity predictors: their incomplete coverage [15]. This is relevant in clinical settings because rejected predictions are associated with specific downstream medical actions. Cost models address these limitations: context is explicitly modeled using cost parameters; incomplete coverage, or rejection rate, is modeled through the inclusion of an additional term (compare equations (3) and (7)). Consequently, there is no averaging over clinical scenarios, and opposite sensitivity/specificity trade-offs give different results.

Here, we use cost models to compare pathogenicity predictors across the clinical space to determine which predictors are preferable in different circumstances. As argued by Drummond and Holte [19], this information can result in valuable insights into the relative values of predictors. It can be helpful to restrict the number of tools to be considered, to define application regions for each tool, etc. However, partitioning a bi-dimensional clinical space (in MISC+REJ) when the number of candidate predictors is arbitrary is a difficult task. Here, we have used the formal properties of cost models to solve this problem for MISC and MISC+REJ.

Applying the cost framework to a set of seventeen pathogenicity predictors with different performances unveils some interesting results. First, context does affect the method of choice (Figs. 1a and 3c), and, consequently, there is not a single preferred method for the whole clinical space. However, the number of selected methods is small, between three and four, which limits the complexity of the tool selection problem. The contrast with the AUC or MCC rankings (Figs. 1b and 3d) shows that cost models expand the view provided by these measures. This is further supported by the comparison between MISC (Figure 1a) and MISC+REJ (Figure 3c) results, which confirms the relevance of considering the rejection rate.

The cost framework developed can also be applied to find the adequate tool(s) for specific genes. Given that the performances of predictors vary between genes [43], we need to estimate the sensitivity, specificity, and coverage of the predictor(s) of interest on a set of variants from the chosen gene. Here, for the TP53 gene, we have retrieved these data from the work of Fortuno et al. [42], who compare a set of predictors for their use in scoring TP53-variants. In Figure 5 (and Supplementary Figures 1c-1d), we look at these predictors using MISC and MISC+REJ. The results are consistent with our previous analyses in that (i) a single predictor is not optimal throughout the whole clinical space, and (ii) the rejection rate of the predictors has a substantial effect on the selected predictors. The latter may be relevant when adapting the ACMG/AMP guidelines because their recommendation for using computational evidence leads to an increased rejection rate. The effect of this situation (Figure 5) opens the way to other alternatives for the rejection area, like using a double threshold system such as the one used in the ATM-adapted ACMG/AMP guidelines [44].

**Figure 5.**
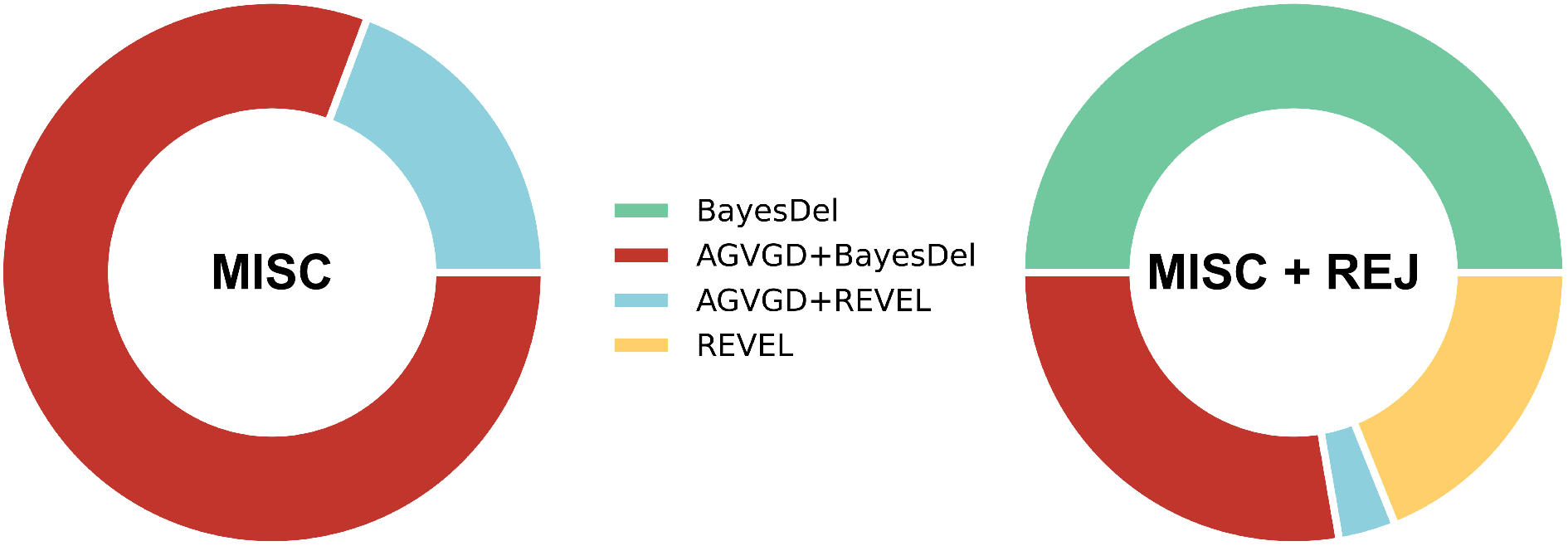
Application of the cost models MISC and MISC+REJ to the predictors used in Fortuno et al. [42] study for TP53 variants. The rings corresponding to MISC (left) and MISC+REJ (right) are divided according to the percentages of cost scenarios corresponding to each predictor. A comparison of both figures shows significant changes in the area occupied by each predictor, primarily due to the presence/absence of the rejection rate in the cost models.

As explained before, we have used the cost framework as a general tool to create insight relevant for the clinical use of pathogenicity predictors. This application does not require knowing the values of the cost parameters rc_0_, rc_1_, and rc_2_. However, we need to know these values to utilize cost models for specific systems, which rarely happens [17]. A possible alternative to overcome this problem is to work with ratios of these parameters because they are easier to estimate by domain experts [17]. It is straightforward to translate this strategy to our models, given the definition of the rci parameters (see Eqs. (3) and (7)) as normalized costs, whose ratios are equal to those of total costs.

Finally, it is worth noting that the methodology presented is independent of the nature of the tools compared (we only need to know their se, sp, and a). Therefore, it can be applied to compare bioinformatics tools in other biomedical/clinical fields like medical imaging [45], diagnosis/prognosis of covid-19 [46], etc., where dozens of classifiers are developed to support healthcare professionals in their work.

## Key Points

- We introduce the use of cost models for the assessment of pathogenicity predictors taking into account clinical context
- We have developed a search strategy for each cost model presented so that multiple pathogenicity predictors can be compared simultaneously across the clinical space
- We illustrate the applicability of the framework developed in a selected set of seventeen pathogenicity predictors
- Our results show that, in addition to the role of context, modeling the two components of the prediction process (misclassification errors and rejection rate) has an important effect on the selection of predictors

## Acknowledgements

The authors acknowledge comments on the work from members of the Pirepred european consortium.

## References

1. Lázaro C, Lerner-Ellis J, Spurdle A. Clinical DNA Variant Interpretation. 2021;

2. Starita LM, Ahituv N, Dunham MJ, et al. Variant Interpretation: Functional Assays to the Rescue. Am. J. Hum. Genet. 2017; 101:315–325

3. Shendure J, Findlay GM, Snyder MW. Genomic Medicine–Progress, Pitfalls, and Promise. Cell 2019; 177:45–57

4. Berrios C, Hurley EA, Willig L, et al. Challenges in genetic testing: clinician variant interpretation processes and the impact on clinical care. Genet. Med. 2021; 1–11

5. Özkan S, Padilla N, Moles-Fernández A, et al. The computational approach to variant interpretation: principles, results, and applicability. Clin. DNA Var. Interpret. Theory Pract. 2021; 89–119

6. Richards S, Aziz N, Bale S, et al. Standards and guidelines for the interpretation of sequence variants: A joint consensus recommendation of the American College of Medical Genetics and Genomics and the Association for Molecular Pathology. Genet. Med. 2015; 17:405–424

7. Wagstaff KL. Machine Learning that Matters. Proc. 29 th Int. Conf. Mach. Learn. 2012;

8. Fortuno C, Lee K, Olivier M, et al. ClinGen TP53 Variant Curation Expert Panel. Specifications of the ACMG/AMP variant interpretation guidelines for germline TP53 variants. Hum. Mutat. 2021; 42:223–236

9. Feliubadaló L, Moles-Fernández A, Santamariña-Pena M, et al. A Collaborative Effort to Define Classification Criteria for ATM Variants in Hereditary Cancer Patients. Clin. Chem. 2021; 67:518–533

10. Hand DJ. Assessing the Performance of Classification Methods. Int. Stat. Rev. 2012; 80:400–414

11. OECD. Health at a Glance 2021: OECD Indicators. 2021;

12. OECD. Health at a Glance: Europe 2020: State of Health in the EU Cycle. 2020;

13. WHO. Pricing of cancer medicines and its impacts. 2018;

14. Mulcahy AW, Whaley CM, Gizaw M, et al. International Prescription Drug Price Comparisons: Current Empirical Estimates and Comparisons with Previous Studies. 2021;

15. Vihinen M. Problems in variation interpretation guidelines and in their implementation in computational tools. Mol. Genet. Genomic Med. 2020; 8:e1206

16. Hanczar B. Performance visualization spaces for classification with rejection option. Pattern Recognit. 2019; 96:106984

17. Adams NM, Hand DJ. Comparing classifiers when the misallocation costs are uncertain. Pattern Recognit. 1999; 32:1139–1147

18. Pepe MS. The Statistical Evaluation of Medical Tests for Classification and Prediction. 2003;

19. Drummond C, Holte RC. Cost curves: An improved method for visualizing classifier performance. Mach. Learn. 2006; 65:95–130

20. Hernández-Orallo J, Flach P, Ferri C. A unified view of performance metrics: Translating threshold choice into expected classification loss. J. Mach. Learn. Res. 2012; 13:2813–2869

21. Hunink MGM, Weinstein MC, Wittenberg E, et al. Decision making in health and medicine: Integrating evidence and values, second edition. Decis. Mak. Heal. Med. Integr. Evid. Values, Second Ed. 2014;

22. de Berg M, Cheong O, van Kreveld M, et al. Computational Geometry: Algorithms and Applications. 2008;

23. Liu X, Wu C, Li C, et al. dbNSFP v3.0: A One-Stop Database of Functional Predictions and Annotations for Human Nonsynonymous and Splice-Site SNVs. Hum. Mutat. 2016; 37:235–241

24. Grimm DG, Azencott C-A, Aicheler F, et al. The Evaluation of Tools Used to Predict the Impact of Missense Variants Is Hindered by Two Types of Circularity. Hum. Mutat. 2015; n/a–n/a

25. Frazer J, Notin P, Dias M, et al. Disease variant prediction with deep generative models of evolutionary data. Nature 2021; 599:91–95

26. Adzhubei IA, Schmidt S, Peshkin L, et al. PolyPhen-2 : prediction of functional effects of human nsSNPs. Nat. Methods 2010; 7:248–249

27. Kumar P, Henikoff S, Ng PC. Predicting the effects of coding non-synonymous variants on protein function using the SIFT algorithm. Nat. Protoc. 2009; 4:1073–1081

28. Rentzsch P, Witten D, Cooper GM, et al. CADD: Predicting the deleteriousness of variants throughout the human genome. Nucleic Acids Res. 2019; 47:D886–D894

29. Schwarz JM, Cooper DN, Schuelke M, et al. MutationTaster2: mutation prediction for the deep-sequencing age. Nat. Methods 2014; 11:361–362

30. Reva B, Antipin Y, Sander C. Predicting the functional impact of protein mutations: Application to cancer genomics. Nucleic Acids Res. 2011; 39:e118

31. Ioannidis NM, Rothstein JH, Pejaver V, et al. REVEL: An Ensemble Method for Predicting the Pathogenicity of Rare Missense Variants. Am. J. Hum. Genet. 2016; 99:877–885

32. Chun S, Fay JC. Identification of deleterious mutations within three human genomes. Genome Res. 2009; 19:1553–1561

33. Choi Y, Sims GE, Murphy S, et al. Predicting the functional effect of amino Acid substitutions and indels. PLoS One 2012; 7:e46688

34. Dong C, Wei P, Jian X, et al. Comparison and integration of deleteriousness prediction methods for nonsynonymous SNVs in whole exome sequencing studies. Hum. Mol. Genet. 2015; 24:2125–2137

35. Carter H, Douville C, Stenson PD, et al. Identifying Mendelian disease genes with the variant effect scoring tool. BMC Genomics 2013; 14 Suppl 3:

36. Pejaver V, Urresti J, Lugo-Martinez J, et al. Inferring the molecular and phenotypic impact of amino acid variants with MutPred2. Nat. Commun. 2020; 11:

37. Niroula A, Urolagin S, Vihinen M. PON-P2: Prediction Method for Fast and Reliable Identification of Harmful Variants. PLoS One 2015; 10:e0117380

38. Bromberg Y, Yachdav G, Rost B. SNAP predicts effect of mutations on protein function. Bioinformatics 2008; 24:2397–2398

39. López-Ferrando V, Gazzo A, De La Cruz X, et al. PMut: A web-based tool for the annotation of pathological variants on proteins, 2017 update. Nucleic Acids Res. 2017; 45:W222–W228

40. Hand DJ. Measuring classifier performance: A coherent alternative to the area under the ROC curve. Mach. Learn. 2009; 77:103–123

41. Baldi P, Brunak S, Chauvin Y, et al. Assessing the accuracy of prediction algorithms for classification: An overview. Bioinformatics 2000; 6:412–424

42. Fortuno C, James PA, Young EL, et al. Improved, ACMG-compliant, in silico prediction of pathogenicity for missense substitutions encoded by TP53 variants. Hum. Mutat. 2018; 39:1061–1069

43. Riera C, Padilla N, de la Cruz X. The Complementarity Between Protein-Specific and General Pathogenicity Predictors for Amino Acid Substitutions. Hum. Mutat. 2016; 37:1013–1024

44. HBOPC VCEP. ClinGen Hereditary Breast, Ovarian and Pancreatic Cancer Expert Panel Specifications to the ACMG/AMP Variant Interpretation Guidelines for ATM Version 1.1. 2022;

45. Liu X, Faes L, Kale AU, et al. A comparison of deep learning performance against health-care professionals in detecting diseases from medical imaging: a systematic review and meta-analysis. Lancet Digit. Heal. 2019; 1:e271–e297

46. Wynants L, Van Calster B, Collins GS, et al. Prediction models for diagnosis and prognosis of covid-19: Systematic review and critical appraisal. BMJ 2020; 369:m1328

