## Appendix 1; Suppl. Figures 1-8; Suppl. Table 1 for "Choosing variant interpretation tools for clinical applications: context matters"

**Appendix 1. Description of the MISC+REJ cost model and proof of the main results underlying the clinical space search algorithm under this model**

Most pathogenicity predictors have an incomplete coverage [1], that is, they are reject classifiers [2]. These classifiers do not produce an output for low-confidence predictions. Therefore, to develop a cost model for pathogenicity predictors, our starting point will be the model developed for reject classifiers [2]. However, this model, in its present form is not completely adequate to healthcare applications because it treats equally misclassification errors due to false positives and false negatives. In healthcare applications this is problematic because the medical consequences of each of these terms may vary significantly. In the following, we present an extended version of the cost model for reject classifiers (to which we will refer as MISC+REJ) in which the misclassification error is split into two terms, corresponding to false positive and false negative errors. We will also present an algorithm for the clinical space search under this cost model.

**NOTE**. In the main body of the article we present a cost model for the split misclassification errors (MISC). All the results about MISC are provided in the article.

In the MISC+REJ model the average misclassification cost of a given predictor, c, is obtained as described in the probabilistic tree diagram in Supplementary Figure 2. Multiplying the probabilities along the branches of the tree and adding the results at the terminating nodes gives:

(1) $c = \alpha\rho(1-s_e)c_0 + \alpha(1-\rho)(1-s_p)c_1 + (1-\alpha)c_2$

where the parameters $c_0$ and $c_1$ are the misclassification costs: $c_0$ is the cost associated to annotating pathogenic variants as benign, and $c_1$ is the cost associated to annotating benign variants as pathogenic. $c_2$ is the cost associated to prediction rejection. In general cost models,

$\rho$ and 1-$\rho$ are the probabilities of the two predicted classes. In our case, $\rho$ will be the proportion of pathogenic variants expected in the deployment context. The value of $\rho$, comprised between 0 and 1, varies with the genome region sequenced (gene panel, whole exome, etc.) and with the population of individuals tested (e.g., the individuals attending a specific hospital unit, ethnic group, etc.). $s_e$ and $s_p$ are the sensitivity and specificity of the pathogenicity predictor; $\alpha$ is its coverage (1-$\alpha$ is the rejection rate). These parameters are estimated testing the predictor in a set of $N_{tot}$ variants as follows:

(1.1) $s_e = \frac{TP}{N_p}$ (1.2) $s_p = \frac{TN}{N_b}$ (1.3) $\alpha = \frac{N}{N_{tot}}$

where N ($N \leq N_{tot}$) is the total number of predicted variants; $N_p$, and $N_b$ are the numbers of predicted pathogenic and benign variants. We have that: $N=N_p+N_b$. And $N_{tot}$-N is the number of rejected predictions.

Following Hernández-Orallo et al. [3], instead of c we will use rc, the normalized average cost, which is obtained after dividing both sides of equation (1) by $c_T$ ($=c_0+c_1+c_2$):

$$(2) \qquad rc = \frac{c}{c_T} = \alpha\rho(1-s_e)rc_0 + \alpha(1-\rho)(1-s_p)rc_1 + (1-\alpha)rc_2$$

where $rc_i=c_i/c_T$ (i=0,2) are comprised between 0 and 1, and $rc_0+rc_1+rc_2=1$.

We can reduce the number of parameters in rc replacing $rc_2$ by $1-rc_0-rc_1$ in equation (2). We obtain, after some reordering:

$$(3) \qquad rc = [\alpha\rho(1-s_e)+\alpha-1]rc_0+[\alpha(1-\rho)(1-s_p)+\alpha-1]rc_1+1-\alpha$$

rc is defined over a triangular region **T** in the $rc_0$-$rc_1$ plane, bounded by the axes $rc_0$, $rc_1$ and the line $rc_0+rc_1=1$ (Figure 3a). **T** is conceptually equivalent to **I** in the MISC case: each point in **T** corresponds to a clinical scenario. We will refer to **T** as 'clinical space' also. However, **I** and **T** differ in that the second is two-dimensional; i.e. clinical scenarios are represented by ($rc_0$, $rc_1$) pairs, not by a single value.

Comparing predictors with either c or rc gives the same result because both quantities are monotonically related (see equation (2)). For N predictors, in a clinical scenario defined by a pair ($rc_0$, $rc_1$), the comparison is straightforward: we only have to compute and sort their respective rc values. The method of choice will be the one with the lowest rc; it will be the predictor with the lowest average MISC+REJ cost.

**Section 1. Generalizing predictor comparison to all clinical scenarios**

The goal here is to give a general solution to the problem of comparing predictors across the clinical space, i.e., for all the values of the cost parameters. The problem is more difficult for MISC+REJ than for MISC because we have now two parameters, $rc_0$ and $rc_1$, instead of one. That is, we have to divide a two-dimensional figure, instead of a one-dimensional interval. To illustrate the procedure, we first describe the case of two predictors (N=2) and then we extend the idea to an arbitrary number N of predictors.

Let $M_i$ and $M_j$ be two pathogenicity predictors, and $rc(M_i)$ and $rc(M_j)$ be their respective rc's. We seek a division of **T** into two regions: $r_i$, where $M_i$ is preferable to $M_j$ ($rc(M_i) < rc(M_j)$), and $r_j$, where the opposite is the case ($rc(M_i) > rc(M_j)$). The boundary between $r_i$ and $r_j$ is defined by the equation $rc(M_i)=rc(M_j)$, which, using equation (3) for $rc(M_i)$ and $rc(M_j)$, gives:

$$(4)\quad \{\rho[\alpha_i(1-s_{e,i})-\alpha_j(1-s_{e,j})]+\alpha_i-\alpha_j\}rc_0+\{(1-\rho)[\alpha_i(1-s_{p,i})-\alpha_j(1-s_{p,j})]+\alpha_i-\alpha_j\}rc_1+\alpha_j-\alpha_i = 0$$

where $s_{e,k}$, $s_{p,k}$ and $\alpha_k$ are the sensitivity, specificity and coverage of predictor $M_k$ (k=i,j). Equation (4) shows that the boundary sought is a line (Figure 3a), which we will call $l_{ij}$, in the $rc_0$-$rc_1$ plane.

When $l_{ij}$ crosses **T,** it divides it into two convex polygons (Figure 3a), corresponding to the $r_i$ and $r_j$ regions. If $l_{ij}$ does not cross **T,** then only one of the two methods will have the lowest rc in all **T** points.

From Equation (4), we see that $l_{ij}$ depends on ρ; consequently, different values of this parameter may change $r_i$ and $r_j$ (Supplementary Figure 3). In this appendix we concentrate on the problem of dividing **T** when more than two predictors are available, keeping the value of ρ fixed. In the main body of the paper, we describe how different values of ρ affect the resulting division.

To generalize the comparison to more than two predictors ($M_i$, i=1,N; N>2), we need a procedure that divides **T** into m regions, {$r_k$, k=1,m}, such that only one method per region has the lowest rc. (Note that m ≤ N, since there may be predictors that are never better than the others). The next two sections are devoted to describing this procedure, first in geometric (Section 2) and then in computational (Section 3) terms.

### Section 2. Dividing T into a set of regions {$r_k$, k=1,m} in which only one predictor per region has the lowest rc

Note. In our proofs we use several results about convex polygons that can be easily found in the books of Lee [4] and of Yaglom and Botyanskii [5]. The most important ones are explicitly cited.

Here, we address, in geometric terms, the problem of finding the {$r_k$, k=1,m} regions in which only one predictor per region has the lowest rc. The starting point is the set of N predictors that we want to compare. Previously, we have seen that there is a line associated to each pairwise comparison (Equation (4)). Therefore, after doing all the possible M=N.(N-1)/2 comparisons between predictors, we obtain a set $L_N=\{l_{ij}, i=1,N-1 \text{ and } j=i+1,N\}$ of lines. A first important result (Proposition 1) is that these lines cut **T**, producing a division of this triangle into a set $P_N$ of convex polygons that will be key to finding the $r_k$ regions.

***Proposition 1***. Let $N \in \mathbb{N}$ be an arbitrary number of methods and let $L_N=\{l_{ij}, i=1,N-1$ and $j=i+1,N\}$ be the set of lines resulting from all the pairwise comparisons between the methods, using rc. These lines cut **T** into a set $P_N=\{p_l\}$ of convex polygons.

*Proof. By induction.*

Base case. For N=2, there is only one line in $L_N$, since there is only one comparison between two methods. When the line contains no interior point of **T**, either because it does not intersect with **T**, or because it is a supporting line of it, $P_2$ will have a single element, **T**, which is convex because it is a triangle. If the line contains at least a point interior to **T**, then it will cut **T** in exactly two points [5]. The line segment uniting these two points is a chord of the polygon [4] and, by the 'Polygon Splitting Theorem' [4], divides **T** into two convex polygons.

Induction step. Here we show that if the proposition is true for N, then it is true for N+1. That is, we want to show that if $L_N$ divides **T** into a set of convex polygons $P_N$, then $L_{N+1}$ divides **T** into a new set of convex polygons that we will call $P_{N+1}$.

We know that the set of lines resulting from the comparison of N+1 methods, $L_{N+1}$, will contain the lines corresponding to the comparisons between the first N methods, $L_N$, and between these N methods and an additional (N+1)th method, $\{l_{i,N+1}\}_{i=1,N}$, that is:

(5) $L_{N+1} = L_N \cup \{l_{i,N+1}, i=1, N\}$

Cutting **T** with the lines in $L_{N+1}$ is equivalent to cutting it with the lines in $L_N$ and then with those in $\{l_{i,N+1}, i=1, N\}$ since order is irrelevant to the final result. Therefore, $P_{N+1}$ will be the result of cutting the polygons in $P_N$ with the lines in $\{l_{i,N+1}, i=1, N\}$. When we cut $P_N$ with the first line, $l_{1,N+1}$, we create a new division of **T** in which each of the polygons split by $l_{1,N+1}$ will be replaced by two children polygons (i.e., no polygon traversed by a line remains in the new division of **T**). Next, we will repeat this process for the remaining lines in $\{l_{i,N+1}, i=1, N\}$ until we obtain $P_{N+1}$. At the end of each step, the division of **T** will be constituted by the set of $P_N$ polygons unaffected by the $l_{i,N+1}$ line (these polygons are convex because the proposition is true for N), and by the children of the affected polygons. Given that the affected polygons are convex (again because the proposition is true for N), the children will also be convex, by the 'Polygon Splitting Theorem' [4]. Therefore, at the end of each step, the resulting division of **T** will be constituted by a set of convex polygons and, consequently, $P_{N+1}$, which is obtained at the end of the final step, will be constituted by convex polygons only. QED.

The polygons in $P_N$ have several characteristics that are relevant for the computational algorithm used to list them (described in Section 3.3). First, their edges are noncollinear line segments that belong either to the $L_N$ lines or to the three segments defining **T.** Second, their vertices can be: the **T** vertices, the intersection points between the $L_N$ lines, and the intersection points between these lines and the triangle edges. Third, for each $L_N$ line, the segment delimited by the intersection points of the line with the triangle is formed by a concatenation of edges from $P_N$ polygons (Supplementary Figure 4). And fourth, the same happens for the three segments defining **T,** which are formed by a concatenation of edges from $P_N$ polygons.

The $P_N$ polygons also satisfy the following lemma.

***Lemma 1***. Let p be a polygon from $P_N$. Then none of its interior points belong to another polygon q $\in P_N$.

*Proof. By contradiction.* Let us assume that there exists a polygon p $\in P_N$ such that one of its interior points belongs to q $\in P_N$. This point will belong to one of the edges of q. The line from $L_N$ containing this edge will cut p at two points[5], that is, it will traverse p. This is in contradiction with the procedure utilized to generate $P_N$, in which any polygon traversed by a line from $L_N$ is removed from the polygon list and replaced by the two children polygons. Therefore, p does not exist. QED.

Finally, we will show a key property of the $P_N$ polygons, used to build the regions $r_k$.

***Proposition 2***. Let $P_N$ be the set of convex polygons obtained after dividing **T** using $L_N$, the set of lines associated to the pair comparisons between N methods. For each polygon $p \in P_N$, the lowest rc value at all its interior points always corresponds to the same method.

*Proof. By contradiction*. Let us assume that the proposition is not true. That is, that there exists a polygon $p \in P_N$ with an interior point m such that $M_i$, the method with the lowest rc value at m, is different from $M_j$, the method with the lowest rc value at the remaining interior points of p. Let us consider n, one of these remaining interior points. Then, according to our assumption, the rc values of $M_i$ and $M_j$ at m ($rc_m(M_i)$ and $rc_m(M_j)$, respectively) and at n ($rc_n(M_i)$ and $rc_n(M_j)$, respectively) satisfy the following inequalities: $rc_m(M_i)<rc_m(M_j)$ and $rc_n(M_i)>rc_n(M_j)$, respectively. This is in contradiction with the fact that the boundary line between $M_i$ and $M_j$ does not pass between m and n, because during the construction of $P_N$ (Proposition 1) any polygon traversed by a line from $L_N$ is removed from $P_N$ and replaced by the resulting children polygons. QED.

We will say that a method $M_k$ is associated to a polygon p, when $M_k$ has the lowest rc value for the interior of p. Note, that $M_k$ will also have the lowest rc value at the line segments defining p.

The polygons in $P_N$ do not necessarily coincide with the $r_k$ regions but can be used to obtain using the following procedure.

**Step 1**. **Find the method associated to each polygon.** For each polygon in $P_N$ we apply the next three steps:

**Step 1.1**. Compute the average of its vertices, which is a point belonging to the interior of the polygon because the polygon is convex.

**Step 1.2.** Compute the rc value, at this average point, for each of the N methods.

**Step 1.3.** Sort the N methods according to their rc's at the average point and choose the method with the lowest rc value. By Proposition 2, this method prevails (has the lowest rc) at all the points interior to the polygon considered. This will be the method associated to the polygon.

**Step 2. Obtaining the {$r_k$, k=1,m} regions.** Each region $r_k$ (Supplementary Figure 5) is obtained as the union of all polygons associated to the same method, $M_k$:

(6) $r_k = \cup_{i\in\Omega_k} p_i$

where $\Omega_k$ are the indexes of the $p_i$ polygons in $P_N$ for which $M_k$ has the lowest rc value. We will say that $M_k$ is the method associated to the region $r_k$.

By application of Proposition 2 to the polygons associated to $M_k$, we know that there is only one method associated to each $r_k$. By the same reason, we know that no other region contains a point for which $M_k$ is the method with the lowest rc. That is, by construction, there are no two regions with the same associated method; therefore, $\{r_k, k=1,m\}$ is the set we are looking for.

Note. For clarity purposes, in this work we do not explicitly treat the fact that, at the boundary between adjacent $r_k$ regions, two methods have the same rc value. This fact has no impact on the results presented, neither in the finding of the $r_k$ regions, nor in computing their surfaces, etc.

### Section 3. Computational obtention of the $P_N$ polygons using an adapted Breadth First Search (BFS)

As we have seen in the previous section, once we know the polygons in $P_N$ it is trivial to obtain the $r_k$ regions, using (6). Here, we describe how we can obtain these polygons computationally. In particular, we show that the problem of finding a $P_N$ polygon, in terms of its constituting vertices, can be modeled as a graph problem, and solved with an adapted version of the BFS algorithm that we will call aBFS. A python implementation of this procedure, CSP-rej (Clinical Space Partition, rejection), is available at:

https://github.com/ClinicalTranslationalBioinformatics/clinical_space_partition

Relative to the program, we want to mention that the code reproduces the results presented in this work and allows users to explore other combinations of predictors. However, it is known that geometric computations may present robustness problems [6], which in our case can happen for low ρ values or large numbers of predictors. Therefore, we cannot discard the possibility that the program sometimes fails to find the division of **T.** In these cases, the program will issue an error message.

*Preliminary results: computing the set of edges and vertices of the* $P_N$ *polygons*

The first step in our approach is to compute the set of lines $L_N=\{l_{ij}$, i=1,N-1 and j=i+1,N$\}$, applying equation (4) to all possible comparisons between the N methods.

The next step is to build VP and EP, the sets of vertices and edges, respectively, of the $P_N$ polygons. For VP, we first compute the intersection between the lines in $L_N$, keeping only the points falling inside **T.** These points are included in VP. Then, we compute the intersection between the lines in $L_N$ and the boundaries of **T.** The resulting points are added to VP. Finally, we include in VP the three vertices of **T.** The resulting set of vertices is used to obtain EP, which is constituted by all the $\overline{v_i v_j}$ ($v_i$, $v_j \in$ VP) line segments uniting two consecutive vertices in an $L_N$ line or in the lines defining **T.** Note that every element of EP corresponds to the edge of a $P_N$ polygon, as shown in the next lemma.

***Lemma 2***. Any segment in EP is the edge of at least one polygon in $P_N$.

*Proof. By contradiction.* We assume that there exists a segment $\overline{v_i v_j} \in$ EP which is not the edge of any $P_N$ polygon. In the following, we show that the possible options for $\overline{v_i v_j}$ lead to a contradiction.

By construction of EP, $\overline{v_i v_j}$ belongs either to one of the lines in $L_N$, or to one of the three lines defining **T.** In all cases, within **T** these lines are formed by a concatenation of edges from polygons in $P_N$ (Supplementary Figure 4). Therefore, $\overline{v_i v_j}$ will overlap with some of these edges. Two situations are possible. It may happen that $\overline{v_i v_j}$ spreads over two or more edges. In this case, some of the vertices of these edges will be comprised between $v_i$ and $v_j$, in contradiction with the fact that $v_i$ and $v_j$ are consecutive. A second possibility is that $\overline{v_i v_j} \subsetneq \overline{v_k v_l}$, where $\overline{v_k v_l}$ is one of the edges in the line. This in contradiction with the fact that $v_k$ and $v_l$ are consecutive. Consequently, $\overline{v_i v_j}$ must be an edge of a $P_N$ polygon. QED.

Before describing how we obtain the $P_N$ polygons, we need to prove a result about polygons sharing more than one edge that will be used to prove our computational procedure.

***Proposition 3.*** Let $p \in P_N$ be a polygon with two edges $\overline{v_i v_j}$, $\overline{v_i v_k} \in EP$ forming a consecutive angle (Supplementary Figure 6). There exists no other convex polygon q, different from p, with $\overline{v_i v_j}$ and $\overline{v_i v_k}$ among its edges and the remaining edges belonging to EP, and such that none of its interior points belongs to another polygon in $P_N$.

*Proof. By contradiction.* We will assume that q exists and explore the different possibilities that arise, showing that they all lead to contradiction. In particular, we will focus our reasoning on the relative position of the points in p and q outside $\overline{v_i v_j}$ and $\overline{v_i v_k}$. There are three possibilities, considering that both p and q are convex.

If in q these points are all interior to p, then the edges of q joining $v_j$ and $v_k$ will be interior to p (Supplementary Figure 7a). Because, these edges belong to EP, they then necessarily belong to polygons in $P_N$ (by Lemma 2). This is in contradiction with the fact that none of the interior points of p belong to another $P_N$ polygon (by Lemma 1).

If all the points in p outside $\overline{v_i v_j}$ and $\overline{v_i v_k}$ are interior to q (Supplementary Figure 7b) then some interior points of q will belong to a polygon in $P_N$, because $p \in P_N$.

Finally, we reach a similar contradiction when both p and q have interior points from one another (in this case, the points may come from full segments or fragments of segments) (Supplementary Figure 7c). QED.

*Building $P_N$ with a graph-based approach*

In our procedure, the list of polygons in $P_N$ is obtained looping through all the vertices in VP, enumerating the polygons that meet at each vertex. Since a polygon is defined by its list of vertices, our approach will produce this list for each of the meeting polygons. Here, we show that we can model this polygon enumeration problem as a cycle enumeration problem in graph theory.

Our starting point is the unweighted, undirected graph G(V, E), whose set of vertices, V, and edges, E, correspond to VP and EP, respectively. Because the list of vertices of a polygon is formally equivalent to that of a cycle, we can reformulate the original looping through VP elements as a looping through V elements. The lists of vertices of the $P_N$ polygons meeting at a given vertex will now correspond to cycles in G(V, E).

In this search, for each $v_i \in V$, we will use BFS as a shortest cycle generator, keeping only those cycles satisfying the following conditions:

- C1. A cycle cannot have more than one edge corresponding to a segment from the same line. This rule is applied to eliminate those sequences of edges produced by the BFS that do not correspond to convex polygons, according to the edge-line lemma [4]. Also, to avoid that edges from collinear segments are included.
- C2. A cycle cannot have repeated vertices, except the first and the last one, as in polygons all the vertices are different except the first and the last one [4].
- C3. Every edge in E has a counter that is decreased by one each time it is included in a cycle. Once the counter reaches zero, the edge is excluded from future searches. The

starting value of the counter of each E edge depends on the location of its equivalent edge in EP. If the latter belongs to a side of the triangle, the counter will start at 1; otherwise (when it belongs to a line in $L_N$) it will start at 2. This condition guarantees that, for each edge in EP, we enumerate all the polygons sharing it, thus ensuring, together with C4, that our polygon enumeration procedure is exhaustive for $P_N$ elements.

- C4. Every vertex in V has a counter that is decreased by one each time it is included in a cycle. Once the counter reaches zero, the vertex is excluded from future searches. The starting value of the counter of each V vertex depends on the environment of its equivalent vertex in VP. More precisely, it depends on the number and origin of EP edges that include the VP vertex (see Supplementary Figure 8). This condition guarantees that, for each vertex, we enumerate all the polygons sharing it, thus ensuring, together with C3, that our polygon enumeration procedure is exhaustive for $P_N$ elements.
- C5. Once a minimal cycle is found, it is excluded from future searches. This condition is introduced to avoid repetitions in the final list of cycles.
- C6. For every minimal cycle found, we check that the corresponding polygon has no interior points corresponding to vertices in VP. This condition, together with C7, prevents the inclusion in the final list of cycles of convex polygons not belonging to $P_N$ (see Lemma 3 and Proposition 4 below).
- C7. For every minimal cycle found, we compute all the possible chords between the vertices of the corresponding polygon. If any of these chords corresponds to an edge in EP, the polygon is discarded. This condition excludes cycles with a list of vertices in which more than two vertices from the same line are included, thus limiting the chosen cycles to those corresponding to convex polygons (all pairs of non-adjacent edges of

the chosen polygon are semiparallel and, by Theorem 8.7 in Lee, the polygon is convex). Condition C7, together with C6, also prevents the inclusion, in the final list of cycles, of convex polygons not belonging to $P_N$ (see Lemma 3 and Proposition 4 below).

As mentioned above, we call aBFS this combination of BFS and C1-C7 conditions. We will now establish (Lemma 3 and Proposition 4) that a cycle found with aBFS corresponds to a polygon in $P_N$.

***Lemma 3***. Let $c_{SC}$ be a cycle found by aBFS, and constituted by the sequence of edges $\{v_i, v_{i+1}\} \in E$, where i=1,N and $v_1=v_{N+1}$. Then, $p_{sc}$, the corresponding sequence of segments $\overline{v_i v_{i+1}}$ from EP, is a convex polygon and has no interior points from any polygon in $P_N$.

*Proof.* $c_{SC}$ is a cycle whose sequence of edges correspond to a sequence of segments, from EP, characteristic of a polygon [4]: it starts and ends at the same vertex, there are no collinear segments (condition C1), and it has not repeated vertices (condition C2) other than the first and last ones, which are equal.

$p_{sc}$ is convex because of conditions C1 and C7.

$p_{sc}$ has no interior points from any polygon in $P_N$ because of conditions C6 and C7. QED.

***Proposition 4***. Let $c_{SC}$ be a Shortest Cycle identified by aBFS. Then, $p_{SC}$, the polygon corresponding to this cycle, belongs to $P_N$.

*Proof.* First, we know from Lemma 3 that $p_{sc}$ is a convex polygon with no interior points from any polygon in $P_N$. Now, let us select an arbitrary pair of adjacent edges from $p_{sc}$, $\overline{v_i v_j}$ and $\overline{v_i v_k}$.

By construction of EP, $\overline{v_i v_j}$ and $\overline{v_i v_k}$ are edges of polygons in $P_N$ (Lemma 2). Because no line from $L_N$ passes between them (by conditions C6 and C7), $\overline{v_i v_j}$ and $\overline{v_i v_k}$ belong to the same polygon $p \in P_N$. From Proposition 3 we know that p is the only convex polygon with $\overline{v_i v_j}$ and $\overline{v_i v_k}$ among its edges and no interior points from any polygon in $P_N$, therefore, $p_{SC} = p$ and $p_{sc} \in P_N$. QED.

For each vertex $v_i$, aBFS will find the shortest cycles corresponding to all the $\overline{v_i v_j}$, $\overline{v_i v_k}$ pairs forming consecutive angles. This procedure will be repeated for all the vertices in G(V, E), guaranteeing, through the use of counters (conditions C3 and C4 above), that the number of cycles found matches that of expected $P_N$ polygons. By condition C5 and Proposition 4, we know that the shortest cycles found are unique and correspond to $P_N$ polygons, respectively. In summary, exhaustive application of aBFS to the vertices in E will produce the list of polygons in $P_N$.

**References**

1. Vihinen M. Problems in variation interpretation guidelines and in their implementation in computational tools. Mol. Genet. Genomic Med. 2020; 8:e1206

2. Hanczar B. Performance visualization spaces for classification with rejection option. Pattern Recognit. 2019; 96:106984

3. Hernández-Orallo J, Flach P, Ferri C. A unified view of performance metrics: Translating threshold choice into expected classification loss. J. Mach. Learn. Res. 2012; 13:2813–2869

4. Lee JM. Axiomatic Geometry. 2012;

5. Yaglom IM, Boltyanskii VG. Convex Figures. 1961;

6. de Berg M, Cheong O, van Kreveld M, et al. Computational Geometry: Algorithms and Applications. 2008;

7. Fortuno C, James PA, Young EL, et al. Improved, ACMG-compliant, in silico prediction of pathogenicity for missense substitutions encoded by TP53 variants. Hum. Mutat. 2018; 39:1061–1069

### Supplementary Figures

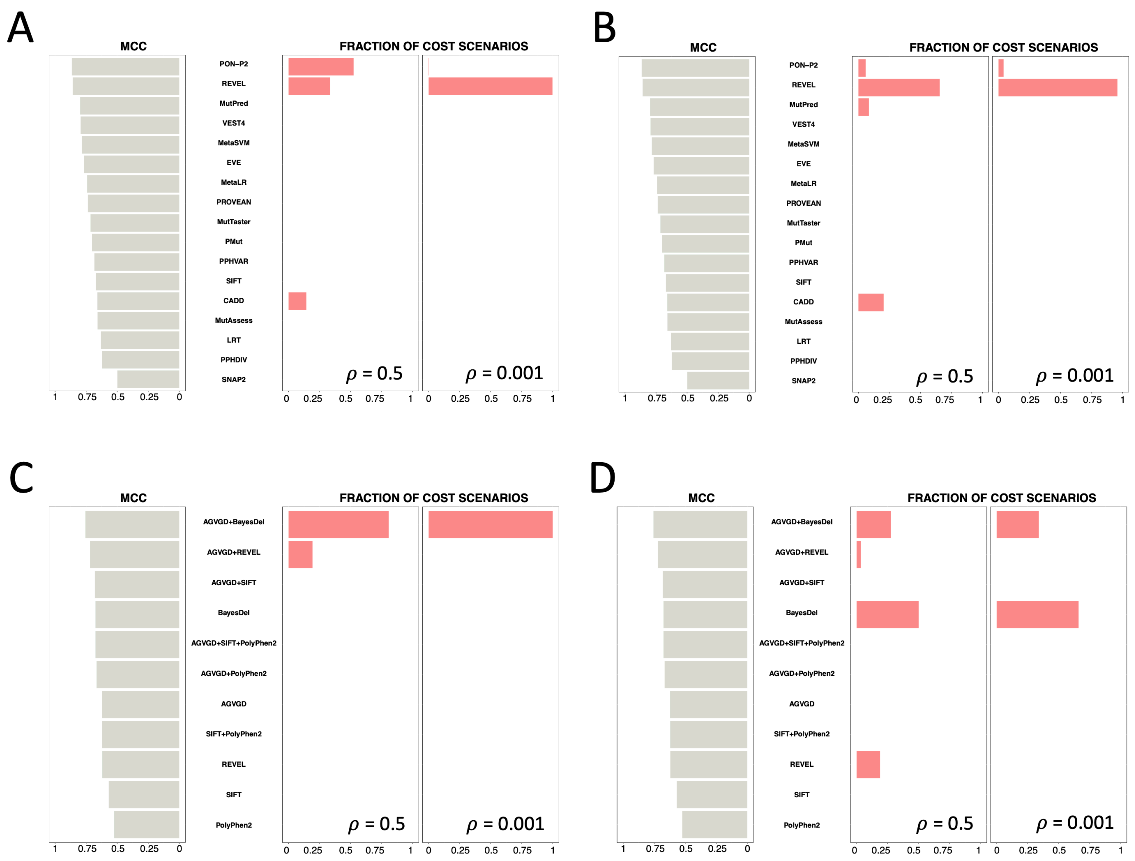


**Supplementary Figure 1. Comparison between the MCC ranking of the seventeen pathogenicity predictors and their corresponding fractions of cost scenarios.** In all the parts of this figure (a-d), pathogenicity predictors are ranked according to their MCC (grey bars, left side) and fraction of cost scenarios for which each predictor is optimal are represented with pink bars, right side. The results are shown for $\rho=0.5$ and $\rho=0.001$). **a**, MISC analysis of the seventeen predictors. **b**, MISC+REJ analysis of the seventeen predictors. These figures are equivalent to those shown in Figs. 1b and 3d, respectively, for AUC. We have reproduced the MCC analysis using data for the TP53 gene and several predictors, retrieved from the work of Fortuno et al. [7]: **c**, MISC analysis, and **d**, MISC+REJ analysis. Here the cost space is restricted to the Li-Fraumeni syndrome. The results confirm an incomplete correspondence between the MCC analysis and the fraction of cost scenarios in which each method predominates.

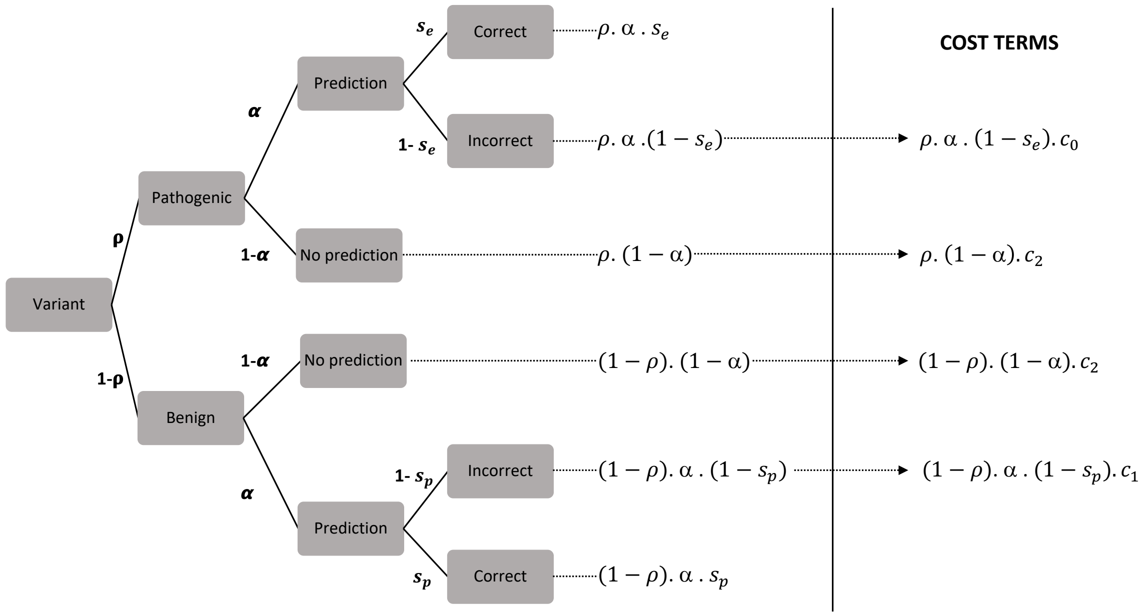


**Supplementary Figure 2. Probabilistic tree diagram underlying the cost-framework presented in this work.** Each branch of the tree corresponds to a different situation in the use of prediction methods, the situation's probability is written by its side. Multiplying probabilities along branches gives the probability of an event that is the combination of different situations. For example, a pathogenic variant can be incorrectly predicted as benign; the probability of this event is: $\rho.\alpha.(1\text{-}s_e)$. In the cost model MISC+REJ each of these events will contribute a term (shown to the right of the vertical line) that, after summation and reordering, will result in equation (1).

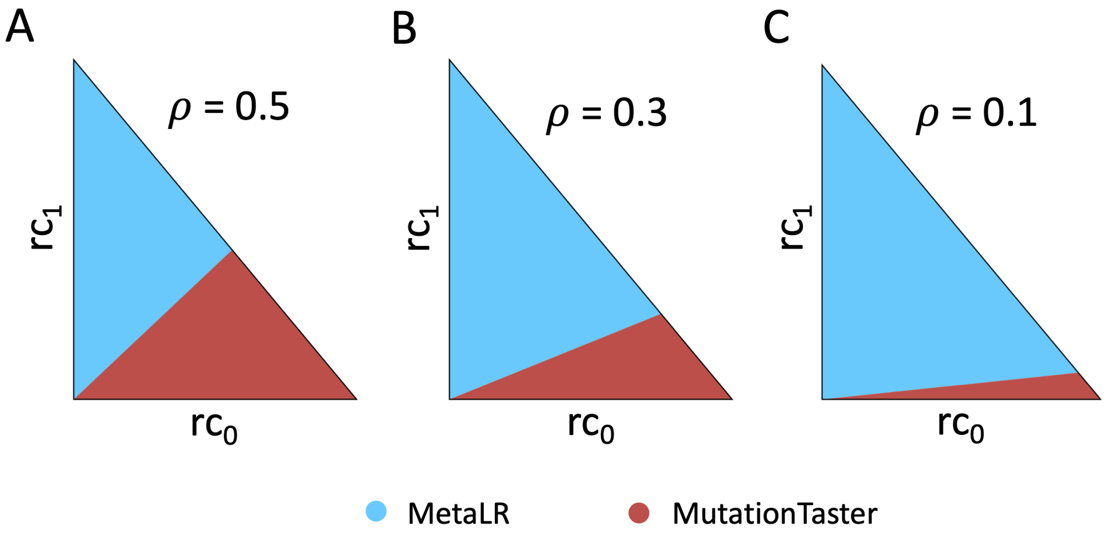


**Supplementary Figure 3. Effect of the fraction of pathogenic variants in the sample (ρ) on the distribution of pathogenicity predictors over the cost domain.** This figure shows how different values of ρ (**a**, ρ = 0.5; **b,** ρ = 0.3; **c**, ρ = 0.1) may substantially alter the size of the cost region assigned to each predictor. A version of this figure for the case of seventeen predictors is presented in Figure 4.

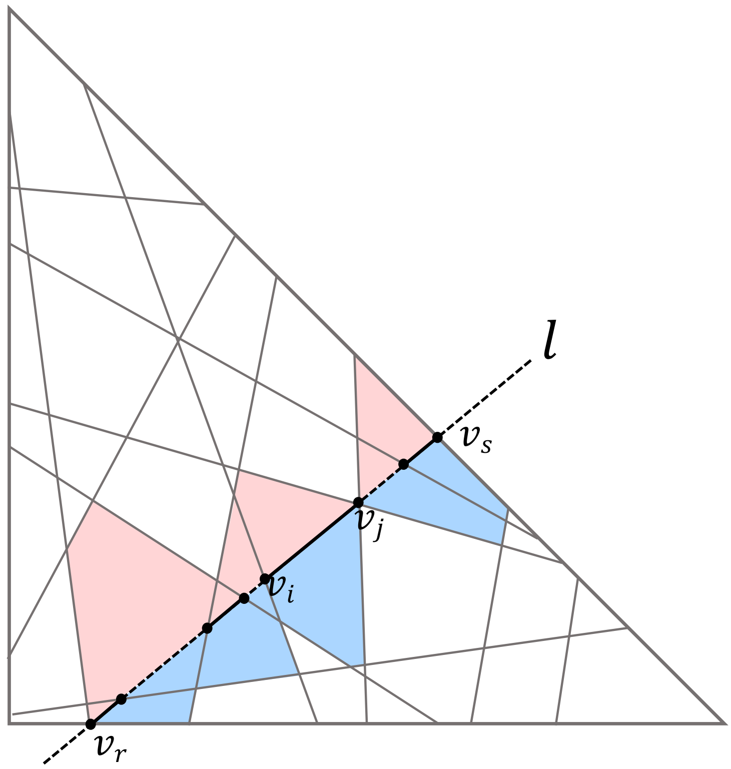


**Supplementary Figure 4. Lines from $L_N$ are constituted by a concatenation of edges from $P_N$ polygons, when crossing the triangle T.** The figure shows how a line $l \in L_N$ is formed by a succession of segments that correspond to edges of $P_N$ polygons (pink/blue), when traversing the triangle **T,** i.e., between the points $v_r$ and $v_s$. As illustrated for $\overline{v_i v_j}$, each segment is shared by two polygons, one above (pink) and the other below (blue) the segment.

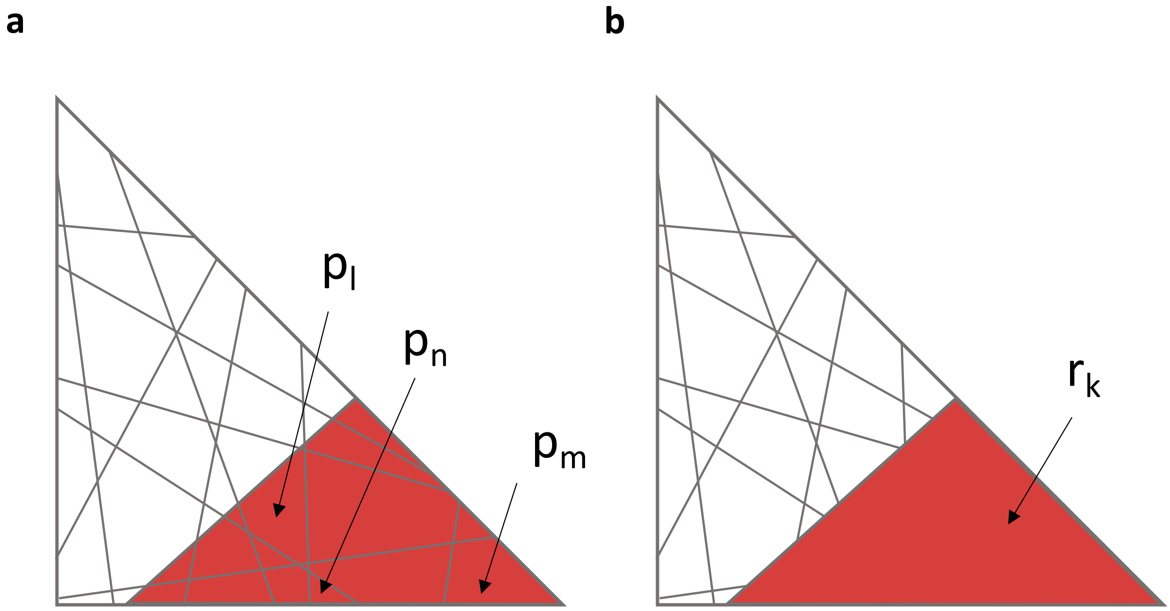


**Supplementary Figure 5. From polygons to regions.** Unification of the polygons in which the same method has the lowest average cost (**a,** shown in red) results in a more simplified view of the regions assigned to each predictor (**b**).

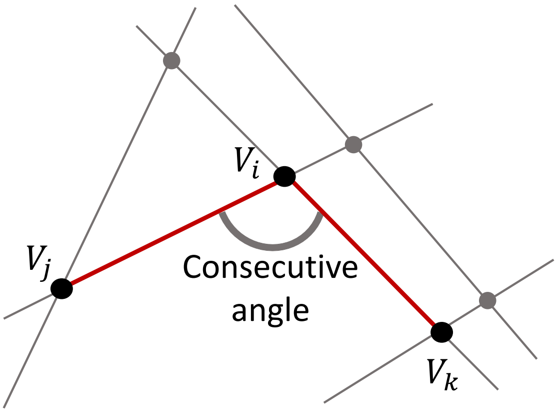


**Supplementary Figure 6. Illustration of the consecutive angle formed by vertices $v_i$, $v_j$, and $v_k$.**

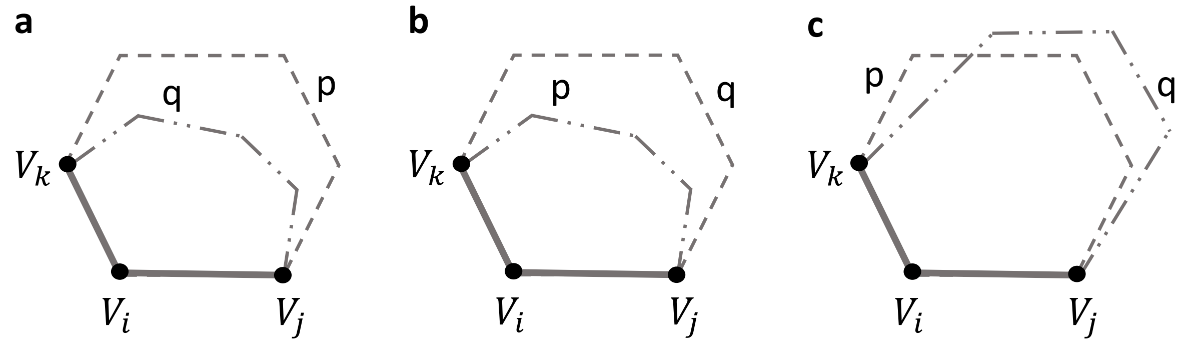


**Supplementary Figure 7. Different situations for the relative location of polygons p and q. a,** all the points in q are interior to p; **b,** all the points in p are interior to q; and **c,** both p and q have interior points from the other.

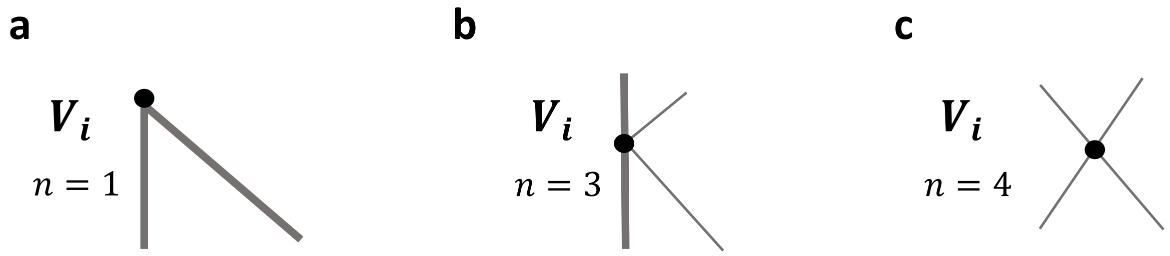


**Supplementary Figure 8. Initial values of the counter of each vertex.** In the three figures, thick grey lines correspond to the edges of the triangle **T** (the clinical space, see Appendix 1, Supplementary Materials), and thin grey lines correspond to the lines dividing **T** and associated with the different pair comparisons between methods. **a,** The vertex is one of the three triangle vertices. **b,** The vertex is the intersection between a triangle edge and the line associated with the comparison between two predictors. **c,** The vertex is the intersection between the lines associated to two different pair comparisons between predictors.

### Supplementary Tables

**Supplementary Table 1. The seventeen pathogenicity predictors used in this work.** We provide the performance parameters required for the cost computations: sensitivity, specificity and coverage. The column Output homogenization shows the correspondence between our pathogenic/benign states and the output of each predictor. We also list the decision cutoff when it was not provided by dbNSFP.

| Predictor | Sensitivity (%) | Specificity (%) | Coverage (%) | Output homogenization & decision cutoff |
|---|---|---|---|---|
| **CADD** | 1 | 0.68 | 1 | P >= 15<br>B < 15 |
| **EVE** | 0.92 | 0.85 | 0.43 | P = Pathogenic<br>B = Benign |
| **LRT** | 0.87 | 0.76 | 0.87 | P = Deleterious<br>B = Neutral |
| **MetaLR** | 0.87 | 0.88 | 0.99 | P = Deleterious<br>B = Tolerated |
| **MetaSVM** | 0.9 | 0.89 | 0.99 | P = Deleterious<br>B = Tolerated |
| **MutPred** | 0.95 | 0.87 | 1 | P >= 0.5<br>B < 0.5 |
| **MutationAssessor** | 0.89 | 0.77 | 0.86 | P = High, Medium<br>B = Low, Neutral |
| **MutationTaster** | 0.98 | 0.74 | 0.99 | P = Disease causing automatic, disease causing<br>B = Polymorphism, polymorphism automatic |
| **PMut** | 0.84 | 0.87 | 0.85 | P = Disease<br>B = Neutral |
| **PON-P2** | 0.96 | 0.92 | 0.46 | P = Pathogenic<br>B = Neutral |
| **PROVEAN** | 0.9 | 0.84 | 0.97 | P = Deleterious<br>B = Neutral |
| **Polyphen2_HDIV** | 0.93 | 0.69 | 0.91 | P = Probably damaging, possibly damaging<br>B = Benign |
| **Polyphen2_HVAR** | 0.9 | 0.78 | 0.91 | P = Probably damaging, possibly damaging<br>B = Benign |
| **REVEL** | 0.92 | 0.94 | 1 | P >= 0.5<br>B < 0.5 |
| **SIFT** | 0.93 | 0.75 | 0.97 | P = Damaging<br>B = Tolerated |
| **SNAP2** | 0.85 | 0.65 | 0.82 | P = Effect<br>B = Neutral |
| **VEST4** | 0.89 | 0.9 | 1 | P >= 0.5<br>B < 0.5 |
